# Differential responses of amphibians and reptiles to land-use change in the biodiversity hotspot of north-eastern Madagascar

**DOI:** 10.1101/2021.03.18.435920

**Authors:** Thio Rosin Fulgence, Dominic Andreas Martin, Romual Randriamanantena, Ronick Botra, Erosiniot Befidimanana, Kristina Osen, Annemarie Wurz, Holger Kreft, Aristide Andrianarimisa, Fanomezana Mihaja Ratsoavina

**Affiliations:** Natural and Environmental Sciences, Regional University Centre of the SAVA Region (CURSA), Antalaha, Madagascar; Zoology and Animal Biodiversity, Faculty of Sciences, University of Antananarivo, Madagascar; Biodiversity, Macroecology and Biogeography, University of Goettingen, Goettingen, Germany; Wyss Academy for Nature, University of Bern, Bern, Switzerland; Sciences of life and Environmental Department, Faculty of Sciences, University of Antsiranana, Madagascar; Tropical Silviculture and Forest Ecology, University of Goettingen, Goettingen, Germany; Agroecology, University of Goettingen, Goettingen, Germany; Centre for Biodiversity and Sustainable Land Use (CBL), University of Goettingen, Goettingen, Germany

**Keywords:** agroecology, agroforestry, community ecology, herpetofauna, human-dominated landscape, land-use history, slash-and-burn cultivation, vanilla, forest dependency

## Abstract

Large expanses of tropical rainforest have been converted into agricultural landscapes cultivated by smallholder farmers. This is also the case in north-eastern Madagascar; a region that retains significant proportions of forest cover despite slash-and-burn hill rice cultivation and vanilla agroforestry expansion. The region is also a global hotspot for herpetofauna diversity, but how amphibians and reptiles are affected by land-use change remains largely unknown. Using a space-for-time study design, we compared species diversity and community composition across seven prevalent land uses: unburned (old-growth forest, forest fragment, and forest-derived vanilla agroforest) and burned (fallow-derived vanilla agroforest, woody fallow, and herbaceous fallow) land-use types, and rice paddy. We conducted six comprehensive, time-standardized searches across at least ten replicates of each land-use type and applied genetic barcoding to confirm species identification. We documented an exceptional diversity of herpetofauna (119 species; 91% endemic). Plot-level amphibian species richness was significantly higher in old-growth forest than in all other land-use types. Plot-level reptile species richness was significantly higher in unburned land-use types compared to burned land-use types. For both amphibians and reptiles, the less-disturbed land-use types showed more uneven communities and the species composition in old-growth forest differed significantly from all other land-use types. Amphibians had higher forest dependency (38% of species occurred exclusively in old-growth forest) than reptiles (26%). Our analyses thus revealed that the two groups respond differently to land-use change: we found less pronounced losses of reptile species richness especially in unburned agricultural habitats, suggesting that reptiles are less susceptible to land-use change than amphibians, possibly due to their ability to cope with hotter and drier microclimates. Overall, old-growth forest harboured a unique diversity, but some species also thrived in vanilla agroforestry systems, especially if these were forest-derived. This highlights the importance of conserving old-growth forests and non-burned land-use types within agricultural landscapes.

## 1. Introduction

Demand for agricultural goods is on the rise due to a growing world population, rising per-capita consumption, and changing diets (Tilman et al. 2011). This situation is leading to both an expansion of croplands into natural areas and an intensification within existing production systems (Tscharntke et al. 2012). Most agricultural expansion in the tropics happens at the expense of forests and leads to an increase in forest fragmentation (Hansen et al. 2020). This expansion affects all aspects of biodiversity: research has documented losses in species richness (Scales and Marsden 2008), shifts in species composition (Newbold et al. 2016), reductions of functional diversity (Matuoka et al. 2020), and erosion of phylogenetic diversity (Li et al. 2020). Importantly, the response of biodiversity to land-use change differs among species groups (Newbold et al. 2014) and world regions (Williams and Newbold 2020), but certain groups and regions are understudied compared to others. While vertebrates are comparatively well studied, we know less about the responses of amphibians and reptiles than we know about mammals and birds (Newbold et al. 2014). Similarly, we know less about what happens in the more biodiverse tropics compared to temperate regions (Gardner et al. 2009). In sum, we know that land-use change is the main driver of biodiversity decline globally (Powers and Jetz 2019), but how the diversity of certain understudied taxa differs between land-use types remains an open question, especially in the tropics, where high land-use pressure and biodiversity coincide (Newbold et al. 2020).

Tropical agricultural landscapes contribute to food security but also provide opportunities for nature conservation (Perfecto and Vandermeer 2010). The quality of the agro-ecosystem for biodiversity conservation depends on the farming systems. Agricultural landscapes dominated by large-scale industrial monocultures have lower conservation value than diverse mosaics of forest fragments, agroforestry systems and more intensively farmed annual crop fields (Mendenhall et al. 2016; Murray and Nowakowski 2021). For example, Ricciardi et al. (2021) reported that small-scale agricultural landscapes host more biodiversity than large-scale ones and De Palma et al. (2015) show that low-intensity agriculture can maintain relatively diverse bee communities. Furthermore, forest patches within tropical small-scale agricultural landscapes increase the variety of ecological niches and benefit species with small ranges in particular (Gray et al. 2016). Besides the value for biodiversity, small-scale land-use mosaics can also provide essential ecosystem services and livelihoods for rural people, making such landscapes work for humans and nature (Kremen and Merenlender 2018). Most research investigating the value of tropical agricultural landscapes for biodiversity and humans was conducted in the Neotropics (Mendenhall et al. 2016) while the conservation value of Afrotropical agricultural landscapes is less understood (Powers et al. 2011). Furthermore, it remains largely unclear how different forms of forest conversion – that is with the use of fire (slash-and-burn shifting cultivation) or without the use of fire (forest degradation and forest-derived agroforestry) – affect amphibian and reptile diversity in resulting land-use types.

Here, we investigate how herpetofauna taxonomic diversity and species composition differ across prevalent land-use types in Madagascar, providing insights into the vulnerability of amphibians and reptiles to various forms of land-use change. Madagascar has lost 44% of forest cover since the 1950s, mainly due to the transformation of natural ecosystems to agricultural lands (Vieilledent et al. 2018). Along with unsustainable extraction rates (Whitehurst et al. 2009), this has resulted in more than half of evaluated Malagasy vertebrate species being at risk of extinction (IUCN 2019). Outstanding levels of endemism (Goodman and Benstead 2005) and ongoing threats qualify Madagascar as a global biodiversity hotspot (Myers et al. 2000). While the forests and protected areas of the island are increasingly well surveyed, the biodiversity in the agricultural landscapes is less known (Irwin et al. 2010). Focusing on amphibians and reptiles is particularly relevant since Madagascar has a diverse and highly endemic herpetofauna: amphibian species diversity is currently estimated at around 370 (AmphibiaWeb 2020) and almost all species are endemic (Goodman and Benstead 2005). Reptile diversity stands at around 440 species (Uetz et al. 2021) with 91% endemism (Goodman and Benstead 2005). Due to cryptic taxonomic complexes, many species still await discovery or description, suggesting that total species richness numbers will increase further (Vieites et al. 2009). Globally, herpetofauna is sensitive to various anthropogenic threats (Vallan 2000) including chytrid fungi, environmental pollution, collection for the pet trade, climate change, and conversion of forest habitat into agricultural lands (Hof et al. 2011). All of these factors are also threatening Malagasy herpetofauna, with deforestation being the top pressure (Cordier et al. 2021).

Our study focuses on north-eastern Madagascar. The area retains more forest cover than other parts of the country (Vieilledent et al. 2018) and is a global priority area for amphibian research (Nori et al. 2018). Besides being known for its remarkable biodiversity, north-eastern Madagascar is also a global centre for vanilla cultivation (Hänke et al. 2018). The price boom of the spice between 2012 and 2019 has triggered an expansion of vanilla agroforests (Llopis et al. 2019), and roughly 80% of rural households in the study region farm vanilla (Hänke et al. 2018). How the expansion of vanilla agroforestry is impacting biodiversity is not well known, despite that the majority of global vanilla cultivation takes place across three biodiversity hotspots (Madagascar, Indonesia and Mexico; Myers et al. 2000, FAO 2020). Investigating how vanilla agroforestry compares to other land-uses in terms of biodiversity conservation thus represents a research gap. Besides farming vanilla, the rural population in north-eastern Madagascar also practices slash-and-burn shifting cultivation for hill rice production. Valleys and plains in the study region are commonly occupied by irrigated rice paddies, forming the backbone of staple crop supply. We thus compared the taxonomic diversity and community composition of amphibians and reptiles between six land-use types within the small-scale agricultural landscape and compared this to continuous old-growth forest inside Marojejy National Park. Within the small-scale agricultural landscape, we sampled vanilla agroforest of contrasting land-use history (forest- and fallow-derived vanilla agroforests), as well as forest fragment, herbaceous fallow, woody fallow and rice paddy. Following Cordier et al. (2021) and Gonzalez and González-Trujillo (2021), we hypothesised that taxonomic diversity would be highest in old-growth forest (for both amphibians and reptiles) and expected losses and shifts in species composition after conversion, particularly in those cases where land-use conversion involves the use of fire for slash-and-burn shifting cultivation.

## 2. Methods

### 2.1. Study region and study design

We conducted our study in the SAVA region in north-eastern Madagascar (Fig. 1A and B) where forests outside protected areas are now highly fragmented (Vieilledent et al. 2018) and the landscape is dominated by smallholder agriculture.

**Figure. 1.**
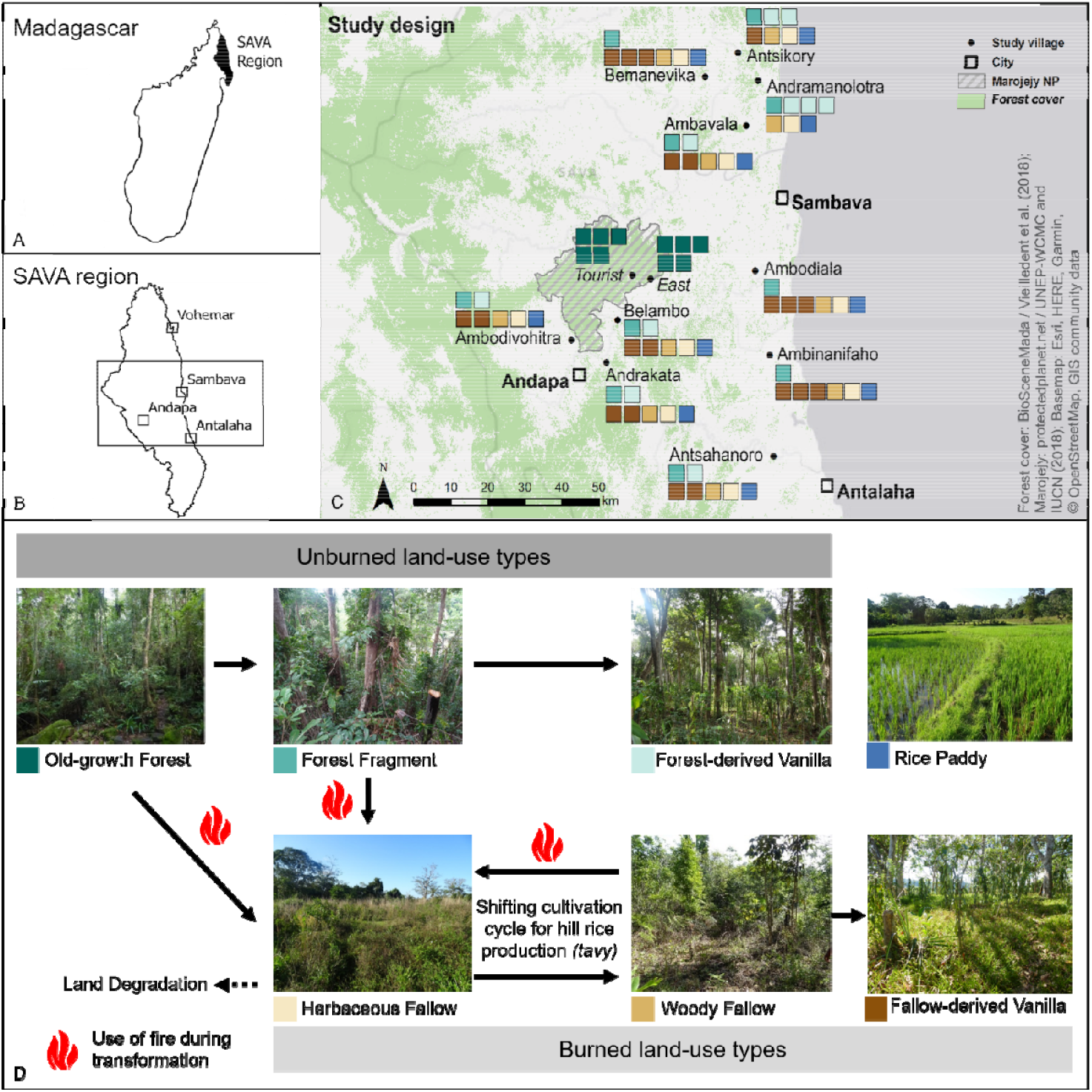
Overview of the study region and the study design. **A** SAVA region in north-eastern Madagascar; **B** study area within SAVA region; **C** study design showing the distribution of 80 plots in ten villages and at two sites inside Marojejy National Park; and **D** overview of the studied land-use types and the typical land-use transformation trajectory from old-growth forest to forest fragments and agricultural land-uses. Panel D modified after Martin et al. (2021) and Panel C modified after Dröge et al. (2021). Rice paddy is not part of the main land-use trajectory.

We collected data on circular plots with a 25 m radius at low to mid-altitude (7-819 m above sea level; mean = 192 m±207 m) surrounding ten villages and in Marojejy National Park, a UNESCO World Heritage Site. In each village, we selected seven plots: three vanilla agroforests (forest-derived and/or fallow-derived vanilla agroforest), one forest fragment, one herbaceous fallow, one woody fallow, and one rice paddy. In the ten villages, we first selected 30 vanilla agroforests along a canopy cover gradient. After consultation with the agroforest owner and a visual confirmation on the plot, we found that twenty vanilla agroforests were fallow-derived while ten agroforests were forest-derived. This approach allowed us to understand the response of amphibians and reptiles to the land-use history of vanilla agroforests. Additionally, we chose ten plots at two sites (five plots each) inside Marojejy National Park. All land-use types, except fallow-derived vanilla agroforests, were replicated ten times. The average minimum distance between one plot and the next closest plot was 719 m± 438 m, while the smallest minimum distance between two plots was 260 m. In total, we surveyed 80 plots (Fig. 1C and D).

### 2.2. Sampled land-use types

We selected 10 *old-growth forest* plots at two sites with five plots each. One of the sites has experienced some selective logging in the past but is now well protected (touristic zone in Manantenina valley; site *Tourist* on Fig. 1), the other site suffers from ongoing occasional selective logging and trapping (Bangoabe; site *East* on Fig. 1), but we chose plots that did not show signs of recent disturbance. The old-growth forest plots are a minimum of 300 m from the National Park boundary. In the study region, isolated *forest fragments* occur around villages and represent remnants of the continuous forest cover that existed in the region prior to the large-scale deforestation that began in the early 20^th^ century (Gade 1996). The ten fragments have not been burned in historic times. The forest fragments are all used for the extraction of timber and non-timber forest products. *Herbaceous fallows* occur after shifting slash-and-burn hill rice cultivation (locally referred to as *tavy*; Styger et al. 2007) and are sometimes used for grazing. The herbaceous fallow plots in this study had last burned at the end of 2016, one year before the onset of data collection in 2017. *Woody fallows* represent successional stages of herbaceous fallows, containing shrubs and small trees. Woody fallows are also occasionally grazed by cows. The woody fallows in our study had last burnt 4-16 years before the onset of data collection in 2017, according to the reports of land-owners. The climbing vanilla orchid (*Vanilla planifolia)* is farmed in agroforestry systems with two distinct land-use histories (following Martin et al. 2020): *forest-derived vanilla agroforests*, where vanilla is directly planted inside the forest after removing understory trees and shrubs and managing tall trees for shade (not involving burning). In *fallow-derived vanilla agroforests*, vanilla is planted on fallow land which resulted from shifting slash-and-burn hill rice cultivation in the past. In these fallow-derived vanilla agroforests, farmers allow natural regeneration of trees or plant trees to provide shade or as support structures for the vanilla vines. Lastly, we studied irrigated *rice paddies* that occur in valley bottoms and flood plains throughout the whole study region. Rice is planted and harvested between one to three times per year. The rice paddies chosen for our study had wider-than-average banks to facilitate movement and sampling within the plots.

### 2.3. Sampling, data collection, and field identification

To collect data in the villages, we organized two sampling campaigns during the driest period of the year (October to December 2017 and late August to December 2018) and one campaign during the wettest period (Mid-January to early April 2018). In Marojejy National Park, we also organized two sampling campaigns during the driest period (late August – early September 2018 and December 2018) and one during the wettest period (February 2019). During each campaign, we visited each plot once during the day (08:00 - 17:00) and once at night (18:30-23:00). Overall, we did six visits per plot, three times during the day and three times at night.

We collected data on amphibian and reptile communities during time-standardized searches (Kadlec et al. 2012). During the survey, we systematically searched each plot in a zig-zag pattern (Kadlec et al. 2012). Each search was standardized to 45 minutes of searching time by two observers. In sum, we thus conducted 270 minutes of searching time on each plot, summing up to 408 hours of searching time across all plots. To detect individuals hiding under rocks, in leaf axils, tree barks, tree holes, leaf-litter, or deadwood, we actively inspected those microhabitats by lifting removable objects to check underneath.

To identify individuals to species level during fieldwork, we used morphological characteristics and referred to keys and visualisations available in ‘A Field Guide to Amphibians and Reptiles of Madagascar’ (Glaw and Vences 2007) and additional literature (Rakotoarison et al. 2017; Ratsoavina et al. 2019). For those individuals which we could not identify with confidence in the field with the available literature, we extracted tissue samples for follow-up DNA analysis and/or collected the specimen. To denote those individuals, we used ‘cf.’ (confer or compare with) and ‘sp. aff.’ (affinis) in conjuncture with the genus name when we found slightly different morphological characters compared to the literature that we used or compared to a given identified species. We used sp. (sp1 or sp2) in conjuncture with the genus name when we could not find any given identified species to compare with the encountered individual. The sp. denotation was thus used to differentiate several unknown species belonging to the same genus and we consider them as morphospecies but used DNA barcodes to confirm species identification (see below). Following the field identification, we resumed the searching time; thus abundance and diversity of amphibians and reptiles were independent of the time spent on the plot. Throughout this manuscript, we refer to each encountered individual as an ‘encounter’ rather than an individual as we cannot exclude the possibility of having encountered the same individual at more than one sampling event.

### 2.4. Species identification with DNA barcodes

We collected muscle or toe clips as tissue samples of individuals in case of non-reliable morphological characteristics based identification. We preserved tissue samples in Eppendorf tubes with 90% ethanol, stored and analysed at the Evolutionary Biology laboratory of Prof. Miguel Vences at the University of Braunschweig, Germany. We preserved specimens in 70% ethanol for amphibians and 70% and deposited them in the collection of the Regional University Centre of the SAVA region (CURSA), Antalaha, Madagascar. We extracted genomic DNA following the standard single-tube salt extraction protocol (Bruford et al. 1992). To do so, we cut small pieces from the collected tissue and proceeded to protein digestion following the protocol of Cacciali et al. (2019). We then passed the extracted DNA to a PCR (Polymerase Chain Reaction) thermocycler, in which we amplified DNA fragments of two mitochondrial genes, “16S” (Vences et al. 2005) and “COI” with the primers 16SAL (5’
s - CGCCTGTTTATCAAAAACAT - 3’) and 16SBH (5’ - CCGGTCTGAACTCAGATCACGT - 3’) from Palumbi et al. (1991). Amplification was set for 45 seconds with 94-96 °C as denaturation, for 45 seconds with 52 °C as hybridization and for 90 seconds with 72°C as elongation. The whole process was repeated for 35 cycles. DNA sequences are generated by sequencer. We first cleaned the obtained sequences with the software CodonCode Aligner version 8.0.2 (Codon Code Corporation, Centerville, MA, US). Then, we processed newly generated sequences for additional quality control with a Blast search (Altschul et al. 1990) which queries similar DNA sequences through GenBank database to provide information on closest taxa or potential contamination. Finally, we built a phylogenetic tree using MEGA-X (Kumar et al. 2018) to reveal clusters of each newly generated sequence, allowing us to reliably identify individuals to species level. We submitted all newly determined sequences to GenBank under accession numbers MZ019097 - MZ019430. In this study, the DNA barcoding differentiated between 43 species of amphibians and 24 species of reptiles. Among them were 12 candidates for new species of which eight were documented for the first time (marked as sp. CaNEW) and four were already found by other researchers (marked as sp. Ca) but have not yet been described. The new species were based on a comparison of obtained DNA sequences to available sequences with the same extraction DNA method conducted in the same lab. In case the DNA barcoding method did not yield results due to short sequences or contaminations, we kept the field identification for those species (for 15 species, eight for amphibians and seven for reptiles).

We attributed the red list status to evaluated species following IUCN (2019). We determined endemicity according to AmphibiaWeb (2020) for amphibians and Uetz et al. (2021) for reptiles. Because most amphibian and reptile genera are completely endemic to Madagascar, we could attribute endemicity also to morphospecies.

### 2.5. Data analysis and visualisation: species diversities and encounters

During the data analysis, we conducted all statistical using R version 3.5.1 (R Core Team 2019) and R package ‘ggplot2’ (Wickham 2016) for visualisation. To compare observed species richness and species encounters on plot-level among land-use types, we computed a generalized linear model with species richness as a response, land-use type as the explanatory variable, and village (respectively old-growth forest site) as a random factor and with Poisson family for count data as distribution. We then ran the *glht* function of the R-package ‘multcomp’ (Hothorn et al. 2008) applying a Tukey all-pair-comparison with Bonferroni correction. We then used the same approach to compare observed species between unburned plots (old-growth forest, forest fragment, forest-derived vanilla) and burned plots (herbaceous fallow, woody fallow, fallow-derived vanilla).

We used the encounter data to compute sample-size-based extrapolation curves (Hsieh et al. 2016) with the ‘iNEXT’ package to assess the diversity (species richness, Shannon, and Simpson diversity) across land-use types using the Hill number framework (Chao et al. 2014). We calculated species richness, Shannon diversity and Simpson diversity (Hsieh et al. 2016). Chao and Jost (2012) define q=0 (0D) as species richness, i.e. the effective number of species in the community, giving equal weight to frequent and infrequent species; q=1 (1D) as Shannon diversity, giving more weight to more frequently observed species and q=2 (2D), as Simpson diversity interpreted as the effective number of abundant species. This approach is now well established given key advantages over traditional Shannon and Simpson diversity indices (Roswell et al. 2021).

To display the total species diversity in each land-use type, we sub-sampled 10 plots of the 20 fallow-derived vanilla agroforests. To do so, we randomly selected one fallow-derived vanilla agroforest from each village. As one of the villages lacks fallow-derived vanilla agroforests (village Andramanolotra, see Fig.1), this resulted in 9 agroforests. We proceeded to select one additional plot from all remaining fallow-derived vanilla agroforests, enabling a fair comparison of total species diversity across 10 plots of each land-use type.

To compare the evenness of communities between land-use types, we additionally plotted the observed and extrapolated diversity across hill numbers for each land-use type, allowing for a direct comparison of slopes between land-use types (Chao and Jost 2015; Roswell et al. 2021).

### 2.6. Data analysis and visualization: species composition

To evaluate the differences in species community composition between land-use types, between burned and unburned plots, and between villages, we used the *metaMDS* function of the R-package ‘vegan’(with 1000 permutations, Oksanen et al. 2020). We used non-metric dimensional scaling (NMDS) of Bray-Curtis dissimilarities to visualize the dissimilarity of species composition in two dimensions. Furthermore, to test the differences between land-use types, we used PERMANOVA (Permutational multivariate analysis of variance) implemented in the *adonis* function of the ‘vegan’ package (Oksanen et al. 2020) and computed pairwise differences using the *pairwise*.*adonis* function with Bonferroni correction of the ‘*pairwise*.*adonis*’ package (Martinez 2020). Since PERMANOVA may confound location and dispersion, we used PERMDISP (Permutational analysis of multivariate dispersions) to see if PERMANOVA results may have been influenced by dispersion differences between groups (Anderson and Walsh 2013). To do so, we used the function *betadisper* and *permutest* and performed pairwise comparisons using the *TukeyHSD* function. To analyse the degree of forest dependency (Rembold et al. 2017), we plotted the proportion of encounters for each species across land-use types. To visualize the forest dependency, we used the *barplot* function of R graphics version 3.5.1.

## 3. Results

### 3.1. Encounters and observed species richness

In total, we made 6215 encounters and observed 119 species of amphibians and reptiles. The 3694 amphibian encounters belong to 58 species, 15 genera, and four families (see SI 1). The most species-rich genera of amphibians are *Boophis* (11 species), *Stumpffia* (10 species) and *Gephyromantis* (eight species). We found all but one species to be endemic according to AmphibiaWeb (2020). Among the observed amphibian species, 22 (37%) could not be identified to species level in the field. The 12 of them were recognized as candidates for new species based on genetic barcoding, another eight of them were counted as morphospecies based on criteria mentioned in the methods section and two were conferred to identified species. Among the encountered amphibian species, seven are listed in the ‘threatened’ category (IUCN 2019). Among the threatened species, we recorded six vulnerable and one endangered species.

The 2521 reptile encounters represent 61 species, 28 genera, and five families (see SI 1). The most species-rich reptile genera were *Phelsuma* (11 species) and *Uroplatus* (six species). We found 83% of reptile species to be endemic (Uetz et al. 2021). Amongst observed reptile species, 15 could not be identified to species level in the field, seven of them were counted as morphospecies and eight were conferred to identified species. Based on the IUCN red list (IUCN 2019), seven encountered reptile species are listed as ‘threatened’. Among the threatened species, we recorded four endangered and one critically endangered species.

Plot-level observed amphibian species richness differed significantly among land-use types (F_6, 73_ =19.59, p-value < 0.001) and when comparing all burned to all unburned land-use types (F_1, 67_ =15.89, p-value < 0.001; Figure SI 6 and table SI 7). A Tukey post-hoc test revealed significant pairwise differences between pairs of land-use types (Fig. 2A and SI 2): Old-growth forests plots had significantly higher observed amphibian species richness while rice paddies plots had significantly lower compared to other land-use types (Fig. 2A). The other land-use types had similar amphibian species richness with no significant differences (see SI 2). Plot level amphibian encounters also differed significantly between land-use types (F_6, 72_ =13.01, p-value < 0.001; Figure SI 9 and table SI 10 and SI 12).

**Figure. 2.**
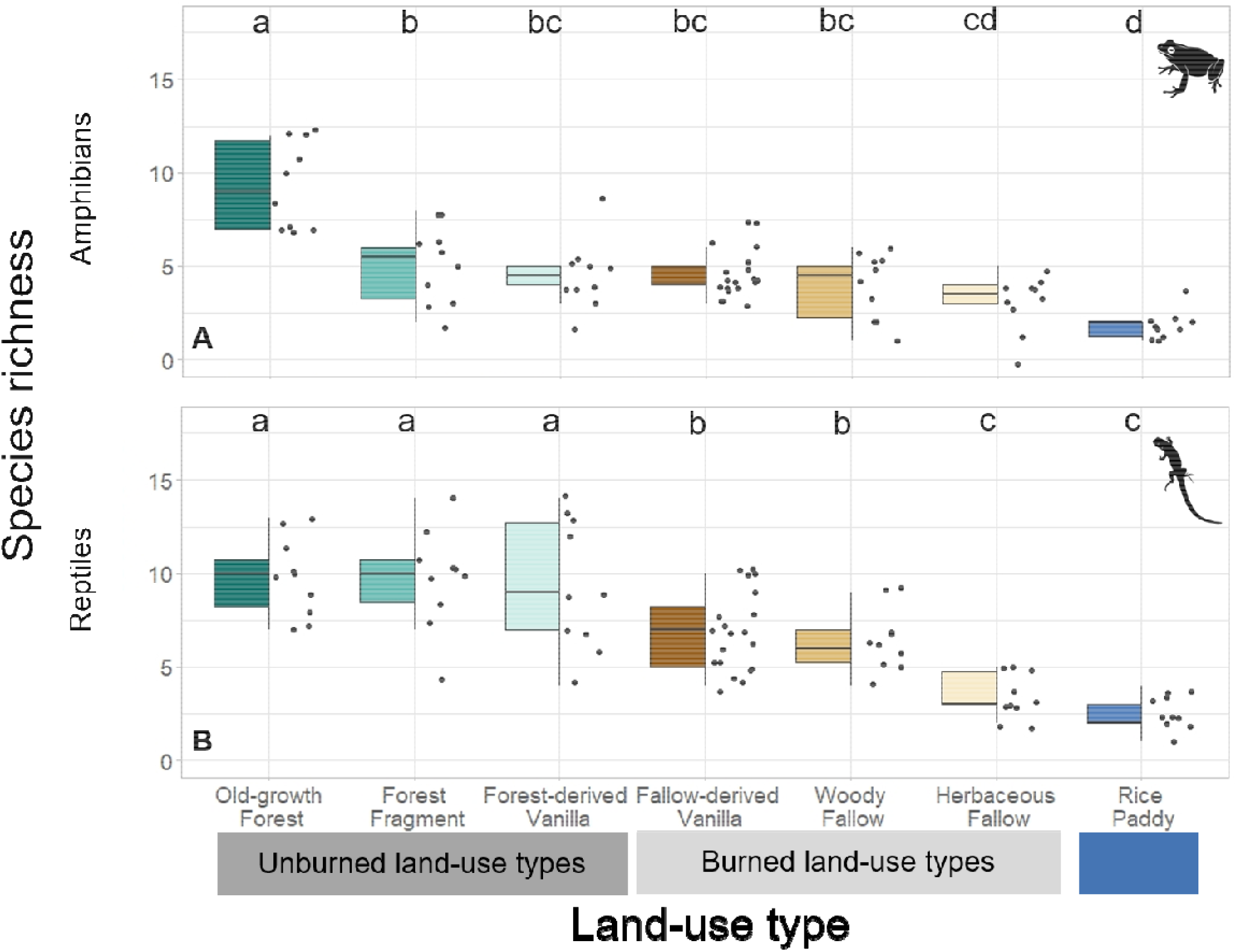
Plot-level observed amphibian (A) and reptile (B) species richness across seven land-use types (replicated 10 times each except fallow-derived vanilla, which is replicated 20 times) in north-eastern Madagascar. The dots represent the species richness per plot in each land-use type. The black horizontal line in the box shows the median. Land-use types with letters in common did not differ significantly based on pairwise comparisons that controlled for inflated false-positive errors using the Tukey HSD approach (Numeric results in SI: amphibians & reptiles SI 2). See icon reference in supplementary information (Free Icons Library).

Plot-level observed reptile species richness also varied significantly among land-use types (F_6, 73_ =18.55, p-value < 0.001) and when comparing all burned to all unburned land-use types (F_1, 67_ =25.60, p-value < 0.001; Figure SI 6 and table SI 8). Tukey post-hoc tests revealed no differences in observed species richness between old-growth forest and forest fragment (p-value = 0.88) and between old-growth forest and forest-derived vanilla agroforest (p-value = 0.77). These three land-use types are recorded with the highest observed species richness, with 10 reptile species on average per plot. Rice paddy had the lowest reptile species richness but there were no significant differences (p-value = 0.19) compared to herbaceous fallow. Reptile species richness was significantly higher in forest-derived vanilla agroforest than in fallow-derived vanilla agroforest (p-value = 0.01; Fig. 2B and SI 2). Plot-level reptile encounters also differed significantly between land-use types (F_6, 73_ =14, p-value < 0.001; Figure SI 9 and table SI 11 and SI 12).

### 3.2. Species accumulation curves, diversity, evenness, and estimated species richness

Encounter-based accumulation curves for amphibians revealed the highest species richness, Shannon and Simpson diversity in old-growth forest and the lowest in rice paddy. In most land-use types, observed amphibian species richness curves flattened off, except for old-growth forest and forest fragment. The overlap of the 95% confidence interval of extrapolated amphibian richness for old-growth forest and forest fragment indicates no differences in species richness. Regarding the accumulated amphibian species diversity for Shannon and Simpson diversity, old-growth forest varied significantly compared to all other land-use types. Shannon and Simpson diversity also did not differ significantly between forest fragment and forest-derived vanilla agroforest, respectively, compared to burned land uses, except to rice paddy (Fig. 3, Table 1).

**Table 1:** Estimated and observed amphibian and reptile species richness (q = 0) for all land-use types and separated per land-use type showing the observed and extrapolated total species richness. Each land-use type is represented by 10 plots; for fallow-derived vanilla, the 10 plots are down-sampled from 20 plots. Extrapolated species richness is based on 5000 encounters and includes the lower and upper 95% confidence interval in italics below (into bracket). See SI for results of q = 1 and q =2 (amphibians and reptiles: table SI 3).

| Land- use type<br>Species Group<br>(measure) | All 80 plots | Old-growth forest | Forest fragment | Forest-derived Vanilla agroforest | Fallow-derived Vanilla agroforest | Woody fallow | Herbaceous fallow | Rice paddy |
| --- | --- | --- | --- | --- | --- | --- | --- | --- |
| **Amphibian**<br>(observed) | 58 | 32 | 26 | 14 | 16 | 8 | 6 | 4 |
| **Amphibian**<br>(extrapolated) | NA | 60.0<br>(20.4 - 99.6) | 46.1<br>(12.4 - 79.8) | 18.0<br>(9.1 - 26.9) | 25.0<br>(7.0 - 42.9) | 9.0<br>(7.4 - 10.6) | 6.5<br>(3.9 - 9.1) | 5.0<br>(4.3 - 5.6) |
| **Reptile**<br>(observed) | 61 | 34 | 30 | 30 | 18 | 19 | 8 | 11 |
| **Reptile**<br>(extrapolated) | NA | 42.0<br>(20.4 - 63.5) | 40.1<br>(10.2 - 69.9) | 35.1<br>(18.2 - 52.4) | 26.9<br>(8.2 - 45.7) | 46.6<br>(11.5 - 81.7) | 8.0<br>(6.8 - 9.2) | 18.9<br>(4.3 - 33.4) |
| **Total**<br>(observed) | 119 | 66 | 56 | 44 | 34 | 27 | 14 | 15 |

**Figure. 3:**
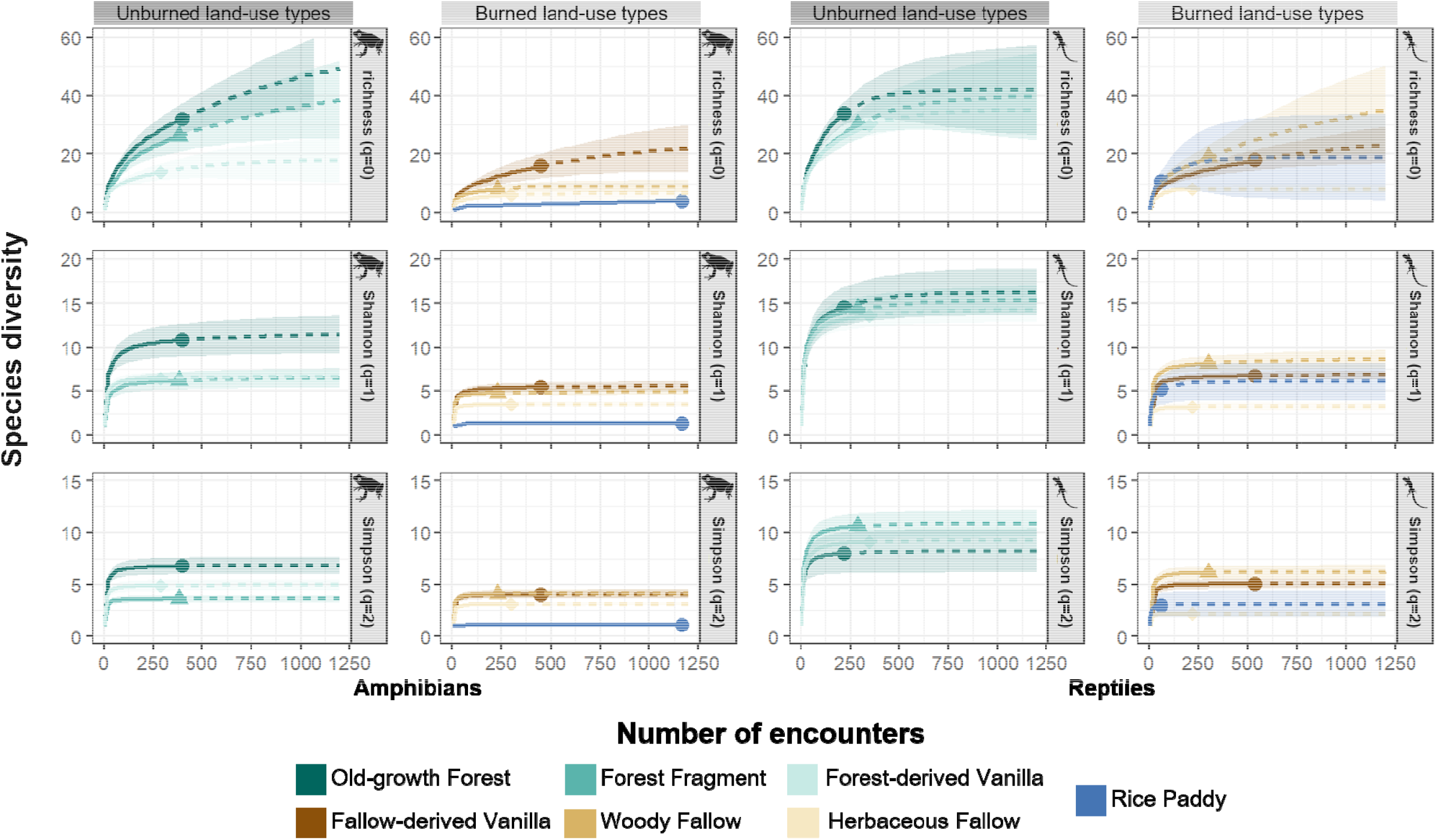
Encounter-based accumulation curves showing interpolated (solid line) and extrapolated (dotted line) diversities for amphibians (first and second column of panel from left to right) and reptiles (third and fourth panel) in north-eastern Madagascar. Unburned (first and third panel) and burned (second and fourth panel) land-use types are separated. The species richness represented by q = 0 (first row of panel from above), Shannon diversity, q=1 (second row of panel) and Simpson diversity q=2 (third row of panel) with 95% confidence intervals (shaded areas) for the amphibian and the reptile data of seven land-use types. The solid dots, triangles and diamonds represent the reference samples, i.e. the observed number of encounters and species richness. See icon reference in supplementary information (Free Icons Library).

Encounter-based accumulation curves for reptiles revealed the highest species richness and Shannon diversity in old-growth forest and the lowest in herbaceous fallow. Observed species richness curves flattened off in all land-use types except woody fallow and fallow-derived vanilla agroforest. We also found an overlap of the 95% confidence intervals of extrapolated species richness as well as Shannon and Simpson diversity within unburned and burned land-use types except for Shannon and Simpson diversity in burned land-use types (Fig. 3, Table 1).

The species diversity dropped more strongly across hill number orders (q=0 to q=2) in amphibians than in reptiles after old-growth forest transformation (Fig. 3, SI 3 & SI 5), highlighting that amphibian communities were more uneven than reptile communities (SI 5). Furthermore, the comparison shows that old-growth forest communities of both amphibians and reptiles were most uneven (SI 5).

### 3.3. Species composition and forest dependency

The composition of amphibian communities differed significantly across land-use types (PERMANOVA: R^2^=0.50, p-value<.001, Df=6), partly driven by differences in group dispersion (PERMDISP: F=4.400, p-value=0.004, Df=6, Table SI 18 – SI 19). Pairwise comparisons showed that amphibian communities in old-growth forest and rice paddy were significantly different from those of other land-use types. Forest fragments differed significantly from fallow-derived vanilla, woody fallow and herbaceous fallow, but no significant differences were observed between forest fragment and forest-derived vanilla. We found no significant differences between forest-derived vanilla, fallow-derived vanilla, woody fallow, and herbaceous fallow for amphibian communities (Fig. 5A, see SI 4). We compared also the amphibian communities between unburned and burned land-use types and found significant differences (PERMANOVA: R^2^=0.193, p-value<.001, Df=1, Fig. SI 13 and Table SI 14), again partly drive by dispersion (PERMDISP: F=34.718, p-value=.001, Df=1, Table SI 16). Similarly, amphibian communities differed between villages and Marojejy National Park communities (PERMANOVA: R^2^= 0.416, p-value<.001, Df=10, Fig. SI 21 and Table SI 22 and SI 24), in this case not driven by differences in group dispersion (PERMDISP: F=1.254, p-value=0.283, Df=11, Table SI 25). Here, communities were very similar between villages (Fig SI 21).

**Figure. 4:**
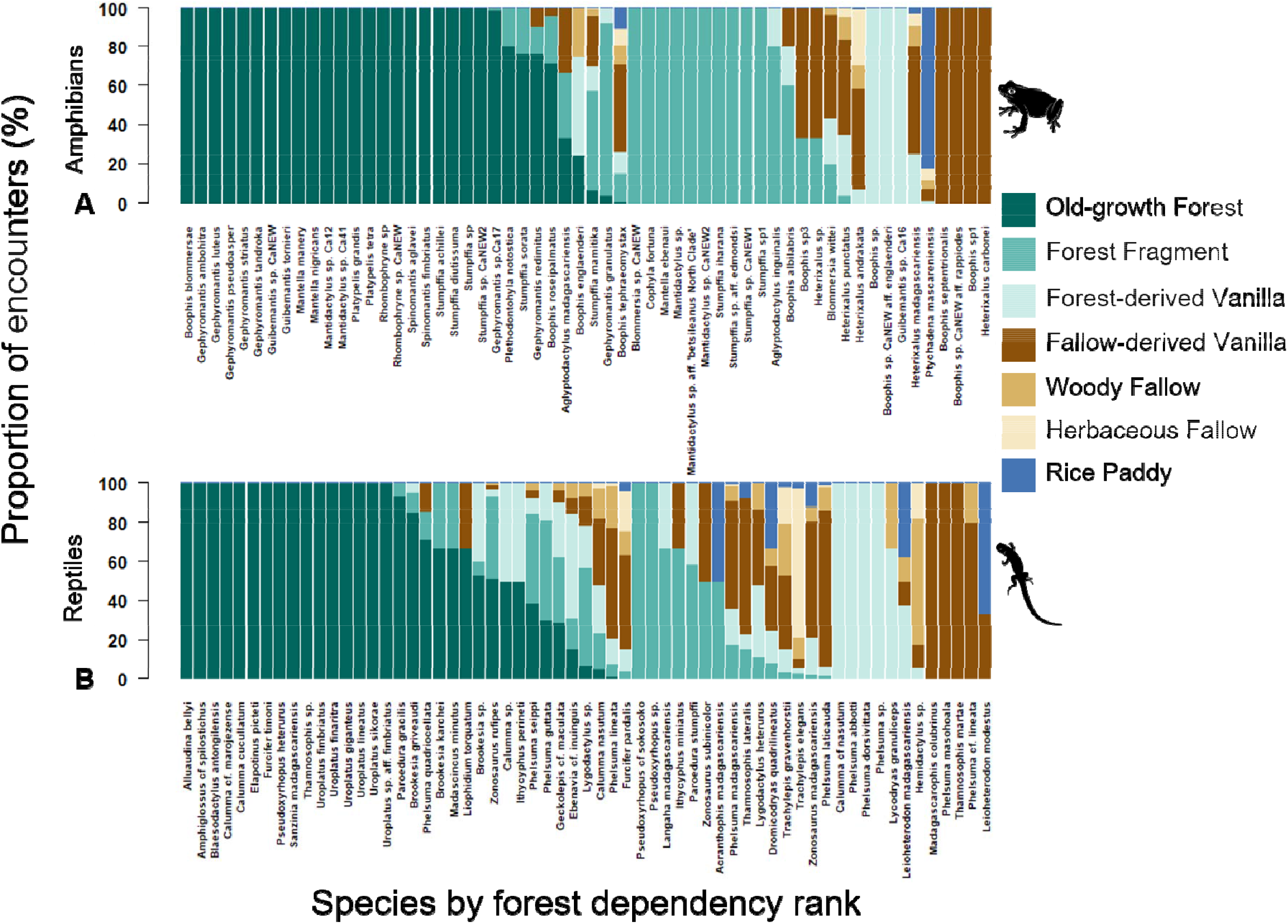
Species composition across seven land-use types in north-eastern Madagascar. All 58 amphibian species with 3694 encounters (A) and all 61 reptile species (B) with 2521 encounters are displayed by forest dependency rank. 38% of amphibian species and 26% of reptile species exclusively occurred in old-growth forest, despite that old-growth forest is only accounting for 12.5% of the total plots. See icon reference in supplementary information (Free Icons Library).

**Figure. 5:**
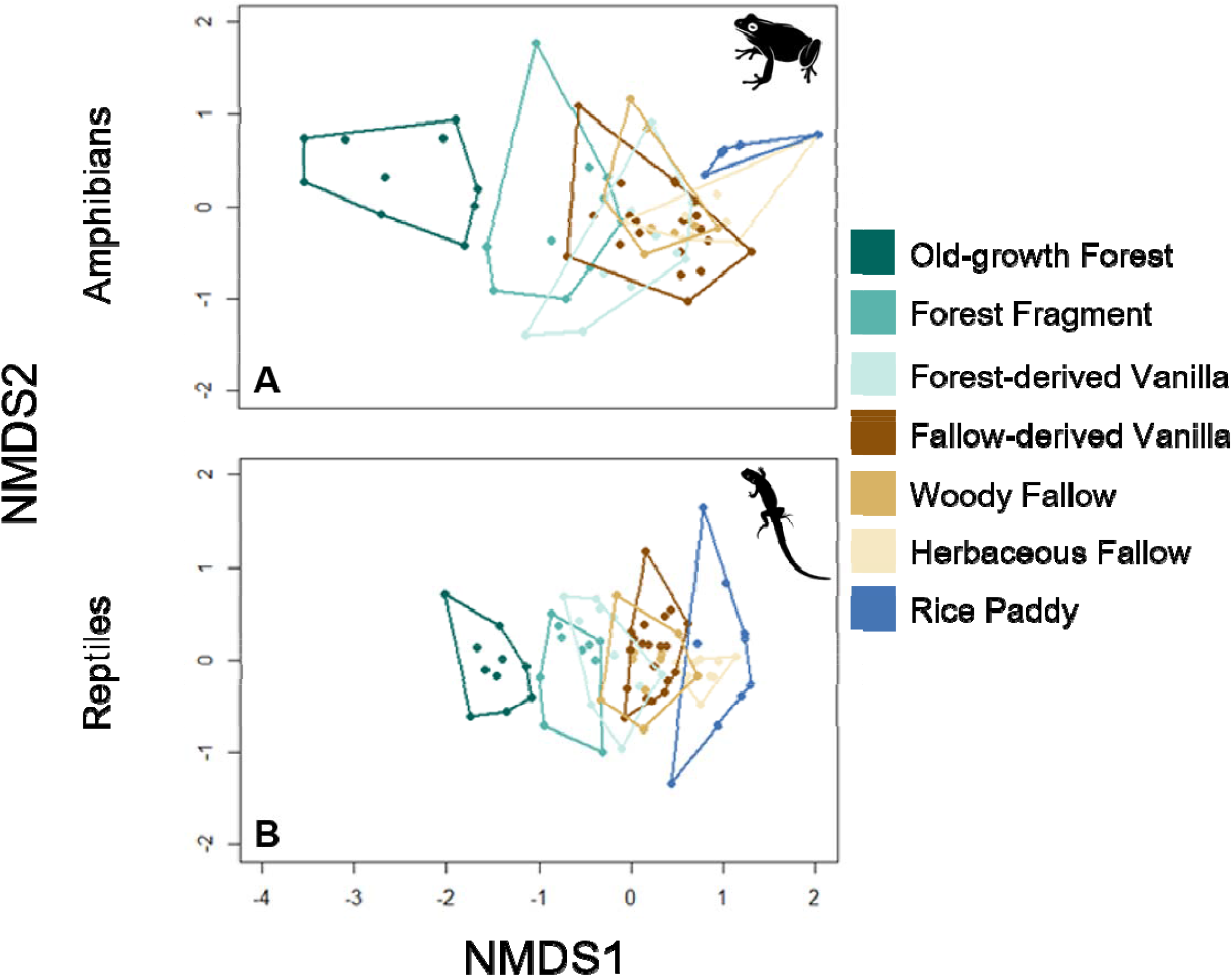
Species composition across seven land-use types in north-eastern Madagascar. Non-metric dimensional scaling (NMDS) showing community dissimilarity of amphibian (A) and reptile (B) communities. Old growth forest shows unique species composition for both amphibians and reptiles See icon reference in supplementary information (Free Icons Library).

Reptile species composition showed significant differences among land-use types (PERMANOVA: R^2^=0.40, p-value<.001, Df=6) and non-significant differences within-group dispersion (PERMDISP: F=1.870, p-value=0.088, Df=6, SI 20) indicating the homogeneity of group dispersion. Pairwise comparisons showed that the reptile communities in old-growth forest and forest fragment were significantly different from all other land-use types. The communities in other land-use types showed no significant differences (Fig. 5B, SI 4). Old-growth forest showed unique communities of reptiles and amphibians. The reptile species composition between unburned and burned land-use types showed significant differences (PERMANOVA: R^2^=0.210, p-value=.001, Df=1, Table SI 15) not driven by differences in group dispersion (PERMDISP: F=3.918, p-value=0.057, Df=1, Table SI 17). The reptile communities between villages and Marojejy National Park showed significant differences (PERMANOVA: R^2^=0.389, p-value<.001, Df=10, Table SI 23 and SI 24) not driven by differences in group dispersion (PERMDISP: F=0.835, p-value=0.626, Df=11, Table SI 26).

Finally, the analyses of forest dependency revealed that 22 amphibian species (38%) were found exclusively in old-growth forest, 11 amphibian species (19%) exclusively in forest fragment, three amphibian species (5%) exclusively in forest-derived vanilla agroforest, and four amphibian species (7%) exclusively in fallow-derived vanilla agroforest (Fig. 4A).

We observed 16 reptile species (26%) exclusively in old-growth forest, three reptile species (5%) exclusively in forest fragment, four reptile species (7%) exclusively in forest-derived vanilla agroforest, and one reptile species (2%) exclusively in fallow-derived vanilla agroforest (Fig. 4B). No species were exclusively found in woody fallow, herbaceous fallow and rice paddy for both groups.

## 4. Discussion

Based on a rigorous study design, extensive field sampling, and DNA-based species identification, we provide a comprehensive assessment of the response of amphibian and reptile species diversity to land-use change in the biodiversity hotspot of north-eastern Madagascar. Overall, we document a highly diverse herpetofauna with 58 amphibian and 61 reptile species and a high proportion of endemic species, 98% for amphibians and 83% for reptiles.

At a plot-level, species richness for both amphibians and reptiles was very high, with up to 12 and 14 species, respectively. For both groups, old-growth forest was significantly different from all other land-use types in terms of total species richness and community composition. Rice paddy for amphibian and herbaceous fallow for reptile harboured the lowest species richness. Reptile species diversity varied significantly between forest-derived and fallow-derived vanilla agroforests whereas amphibian species showed no difference. For woody fallow, herbaceous fallow and rice paddy, no unique species occurred in any of these land-use types. Importantly, we found that amphibians and reptiles responded differently to land-use history. After any kind of old-growth forest conversion, whether through burning or not, amphibian species communities were decimated and showed a high shift in community structure compared to old-growth forest. In reptile communities, the forest alteration through slash-and-burn showed a strong species loss as well, but changes were less pronounced if old-growth forest was transformed to forest fragment or forest-derived vanilla agroforest since these transformations refrain from using fire.

### 4.1. Outstanding diversity of amphibians and reptiles in north-eastern Madagascar

The diversity of amphibians and reptiles documented here was exceptional, both within Madagascar and compared to other tropical biodiversity hotspots. The few available studies on the response of amphibian communities to land-use change in Madagascar reported lower overall values, with 32 and 28 species, respectively (Vallan 2000; Ndriantsoa et al. 2017). Andreone et al. (2000) found 42 amphibian and 23 reptile species in north-eastern Malagasy rainforest. Despite differences in methodology and study sites, our study provides additional evidence that the north-east of Madagascar is one of the most species-rich regions for herpetofauna, as previously suggested (Brown et al. 2016). The documented diversity also exceeds values found in other tropical biodiversity hotspots (Cordier et al. 2021; Murray and Nowakowski 2021) – for example, Urbina-Cardona et al. (2006) found 21 amphibian and 33 reptile species in region of Veracruz, Mexico, and Kurz et al. (2014) found 25 amphibian and 20 reptile species in region of north-eastern Costa Rica (versus 58 amphibian and 61 reptile species in this study). The high species diversity may partly be driven by our extensive sampling effort, in terms of number of plots (total n=80), diversity of different land-use types (seven), and search effort (total of 270 min searching time by two observers per plot), and is likely also influenced by the use of genetic samples for species identification.

### 4.2. Response of amphibian diversity to land-use change

We found a strong negative response of amphibian species richness to any anthropogenic land uses. Old-growth forest had a significantly higher observed amphibian species richness with 38% species exclusively occurring there. Besides, the amphibian species composition is unique compared to all other land-use types, driven by a high share belonging to the Microhylidae family and *Spinomantis, Guibemantis*, and *Mantella* genera (Fig. 4A). Only forest fragments harboured comparably high values to old-growth forest for accumulated amphibian species richness. Subsequently, we infer that amphibians are very sensitive to habitat change. This result is in line with findings from Ndriantsoa et al. (2017) in disturbed habitats of eastern Madagascar which documented strongly impoverished frog communities in secondary vegetation and rice fields compared to forests. Forest fragments are thus valuable for Malagasy amphibians to maintain diversity within the agricultural landscape, yet forest fragments cannot substitute large and continuous old-growth forest (Vallan 2000).

The other land-use types (fallows that form part of the slush-and-burn cycle, vanilla agroforests, and rice paddies) are of minor importance for amphibian conservation, given the low species diversity that consists mainly of common and widely ranged species. Nonetheless, amphibians could play an important functional role in these habitats: abundance of amphibians is high throughout, reflected by high number of encounters, particularly in rice paddies (see Fig. 3B and SI 9). As such, they may be an important food source for other taxa or could provide pest control services in crops (Hocking et al. 2014).

The strong negative response of amphibians to deforestation could be explained by the fact that numerous species rely on moist environments to avoid dehydration (Watling and Braga 2015), especially species living in evergreen forest habitat (Hof et al. 2011). Given that selective logging, forest fragmentation and deforestation severely change microclimate conditions (Ewers and Banks-Leite 2013), many species may struggle to cope. Furthermore, amphibians have different habitat requirements throughout their life cycle stages (Hof et al. 2011; Hocking et al. 2014). Such heterogeneity is often lost in concert with the simplification in forest structure (Wanger et al. 2009), making new habitats unsuitable for many species.

### 4.3. Response of reptile diversity to land-use change

We found a strong effect of land-use history on reptile diversity. Unburned land-use types had a significantly higher average species richness, more uneven communities, more unique species, and a distinct species composition compared to burned land-use types and rice paddy. The distinct species composition in unburned land-use types was caused by an increased abundance compared to other land-use types of species belonging to the Chamaeleonidae family and *Uroplatus* genus (Fig. 4B). This is, to our knowledge, the first study shedding light on the response of reptiles to land-use change in Madagascar’s humid eastern landscape. Studies from the drier southern region (Scott et al. 2006; Gardner et al. 2016; Nopper et al. 2018) show that Malagasy reptile species react less strongly to habitat change than other taxa, especially if diurnal (Nopper et al. 2018).

Interestingly, the less pronounced effects of land-use change on reptiles compared to amphibians in tropical landscapes has also been demonstrated by others (Wanger 2010; Kurz et al. 2014) and was found in a review by Palacios et al. (2013). This may be due to the high thermotolerance in reptiles (Thompson and Donnelly 2018) thanks to a fairly impermeable skin covered with scales. This structure prevents excessive evaporative water loss and adapts reptiles to various microclimates (Watling and Braga 2015). Furthermore, egg development is not heavily impacted by temperature rise, which reduces the size of hatchlings and accelerates incubation (Packard et al. 1982). Reptile lifestyle and traits may thus enable reptiles to adapt to more open, hotter, and drier environments (Morin 2005), making them less vulnerable to land-use change.

### 4.4. Land-use history of vanilla agroforests affects reptiles but not amphibians

In our study, we have separated forest-derived vanilla from fallow-derived vanilla agroforests, thereby explicitly accounting for land-use history (Martin et al. 2020). For amphibians, we found no differences between the two agroforestry systems across metrics. Reptile communities, on the other hand, were significantly more diverse in forest-derived vanilla agroforests on plot level, more species-rich overall, more uneven, and compositionally different compared to fallow-derived vanilla. Reptile communities observed in forest-derived vanilla agroforests are thus more similar to old-growth forest and forest fragment. The reptile communities recorded in fallow-derived vanilla agroforests were comparable to fallow land. Both kinds of agroforest thus resembled the land-use types they were derived from. An increase in accumulated amphibian species richness is also demonstrated from fallow-derived vanilla agroforest over fallow land, which may emphasize the rehabilitation opportunity of fallow land through agroforestry (Osen et al. 2021). These findings are in line with the prediction from a recent study of Martin et al. (2020) which found that the land-use history of agroforestry systems matters for biodiversity.

We further hypothesize, that the strong importance of forest-derived vanilla agroforests for reptiles may be in part driven by leaf-litter depth, which is known to positively influence reptile diversity and abundance (Urbina-Cardona et al. 2006). According to the same study, these microenvironmental variables also showed positive effects on amphibians and thus cannot explain why there are no differences between the two kinds of agroforests for amphibians. Elsewhere, the loss of canopy cover had negative impacts on amphibian and reptile species richness (Scott et al. 2006). Since canopy cover differed significantly between forest-derived vanilla agroforest (higher) and fallow-derived vanilla agroforest (lower) in the same study site (Osen et al. 2021), this is calling for further investigation combining habitat characteristics with species traits (Oliveira et al. 2017) to elucidate the drivers of change in both species groups.

### 4.5. Conservation implications

The strong negative response of amphibians to old-growth forest modification and the high old-growth forest dependency of amphibians and reptiles calls for the effective protection of the last remaining old-growth forests. Additionally, conserving forest fragments within the agricultural landscape will be important to many reptile and amphibian species that are absent from other land-use types. This is particularly important given that numerous species are micro-endemics (Brown et al. 2014), meaning that they may only consist of few fragmented populations (Vieites et al. 2009). Management strategies are thus needed to safeguard the long-term existence of these fragments and to improve the connectivity between fragments. The protection of large continuous forests throughout the region is also important under a changing climate (Hof et al. 2011) and in light of emerging threats, such as the chytrid fungi (Kolby and Skerratt 2015) and the recent spread of the invasive Asian common toad (*Duttaphrynus melanostictus*, Soorae et al. 2020). These conservation needs are underscored by the exceptional diversity of reptiles and amphibians in north-eastern Madagascar as well as by the high proportion of endemic species.

Our findings from the north-eastern Madagascar also have broader relevance to herpetofauna conservation in other tropical biodiversity hotspots. The differential response of amphibians and reptiles to land-use change shows that conservationists should not treat herpetofauna as a homogenous group when devising conservation programs. Furthermore, the distinct differences in reptile communities between plots of contrasting land-use history shows that past land-use should also not be overlooked, suggesting that the maintenance of remnant forest fragments and forest-derived agroforests is important. Lastly, we highlight the importance of unburned compared to burned land-use types in the agricultural matrix for the conservation of reptiles.

## 6. Acknowledgements

We are grateful to all *chef de fokontany*, landowners, and Madagascar National Parks for granting us access to sites and information. We thank Prof. Miguel Vences and his laboratory staff for invaluable support with the DNA barcoding, Saskia Dröge for preparing panel C of Fig. 1, James Herrera for feedback, and Prof. Arne Mooers, Philippe Fernandez-Fournier, and an anonymous reviewer for their constructive reviews. We also thank ‘CodonCode grants’ for offering us a free two-year license under the CodonCode Aligner License Grant Program. We collected data under research permits N°100/17/MEEF/SG/DGF/DSAP/SCB.Re, N°163/17/MEEF/SG/DGF/DSAP/SCB.Re, N°18/18/MEEF/SG/DGF/DSAP/SCB.Re and N°254/18/MEEF/SG/DGF/DSAP/SCB.Re granted by the Ministry of Environment and Sustainable Development, Antananarivo. We transported DNA samples domestically under the transport permit N° 34/17-MEEF/SG/DREEF/SAVA/SRF granted by the Regional Office of the Ministry of Environment and Sustainable Development, Sambava. We exported the samples from Madagascar under CITES permits N°323C-EA05/MG18 and N°180C-EA03/MG19 granted by the Ministry of Environment and Sustainable Development, Antananarivo, and imported the samples to Germany under CITES permits DE-E-03377/18, DE-E-03378/18, DE-E-02422/19, DE-E-02423/19, DE-E-02424/19, and DE-E-02425/19 granted by the German Federal Agency for Nature Conservation, Bonn. This study was financially supported by the Niedersächsisches Vorab of Volkswagen Foundation as part of the research project ‘Diversity Turn in Land Use Science’ (Grant number 11-76251-99-35/13 (ZN3119)) and by the German Academic ExChange Service (DAAD) within the ‘Partnerships for Supporting Biodiversity in Developing Countries’ initiative (Project Nr. 57449386).

## 8. Author’s contributions

TRF, DAM, KO, AW, HK, AA, and FMR designed the study. TRF, EB, RB, and RR collected amphibian and reptile data under the lead of TRF. TRF and DAM analysed the data. TRF and DAM wrote the first manuscript draft. All authors revised the manuscript.

## 9. Data availability

Data are available from Zenodo: http://doi.org/10.5281/zenodo.4548955

