## Supplementary Information for "Differential responses of amphibians and reptiles to land-use change in the biodiversity hotspot of north-eastern Madagascar"

**Authors’ affiliations:**

^1^ Natural and Environmental Sciences, Regional University Centre of the SAVA Region (CURSA), Antalaha, Madagascar

^2^ Zoology and Animal Biodiversity, Faculty of Sciences, University of Antananarivo, Madagascar

^3^ Biodiversity, Macroecology and Biogeography, University of Goettingen, Goettingen, Germany

^4^ Wyss Academy for Nature, University of Bern, Bern, Switzerland

^5^ Sciences of life and Environmental Department, Faculty of Sciences, University of Antsiranana, Madagascar

^6^ Tropical Silviculture and Forest Ecology, University of Goettingen, Goettingen, Germany

^7^ Agroecology, University of Goettingen, Goettingen, Germany

^8^ Centre for Biodiversity and Sustainable Land Use (CBL), University of Goettingen, Goettingen, Germany

**Corresponding author**: Thio Rosin Fulgence; Natural and Environmental Sciences, Regional University Centre of the SAVA Region (CURSA), Antalaha, Madagascar;

Zoology and Animal Biodiversity, Faculty of Sciences, University of Antananarivo, Antananarivo, Madagascar,

Biodiversity, Macroecology and Biogeography, University of Goettingen, Goettingen, Germany

**Table SI 1**: List of encountered amphibian and reptile species per land-use types with IUCN status and endemicity to Madagascar, ‘***cf.***’ and ‘***sp. aff.***’: confer to identified species, ‘***sp.***’: not identified species and shows dissimilarity morphological characters to identified species, ‘***sp. CaNEW***’ (sometimes followed by number, e.g. sp. CaNEW1): Candidate for new species according to DNA barcoding recorded during the current study, ‘***sp. Ca***’ (sometimes followed by a number, e.g. sp. Ca17): Candidate for new species according to DNA barcoding which has already been recorded by other researchers, **LC**: Least Concern, **NA**: Not Applicable, **VU**: Vulnerable, **NE**: Not Evaluated, **EN**: Endangered, **NT**: Near Threatened, **CR**: Critically Endangered, **OGF** (Old-growth forest), **FF** (forest fragment), **VFST** (Forest-derived vanilla agroforest), **VFLW** (Fallow-derived vanilla agroforest), **WF** (woody fallow), **HF** (herbaceous fallow), **RP** (rice paddy).

| **family** | **genus** | **species** | **status** | **endemicity** | **OGF** | **FF** | **VFST** | **VFLW** | **WF** | **HF** | **RP** |
| --- | --- | --- | --- | --- | --- | --- | --- | --- | --- | --- | --- |
| **Amphibian** | | | | | | | | | | | |
| Hyperoliidae | *Heterixalus* | *Heterixalus andrakata* | LC | endemic | 0 | 1 | 1 | 1 | 1 | 1 | 1 |
|  |  | *Heterixalus carbonei* | LC | endemic | 0 | 0 | 0 | 1 | 0 | 0 | 0 |
|  |  | *Heterixalus madagascariensis* | LC | endemic | 0 | 0 | 1 | 1 | 1 | 1 | 1 |
|  |  | *Heterixalus punctatus* | LC | endemic | 0 | 1 | 1 | 1 | 1 | 1 | 0 |
|  |  | *Heterixalus sp.* | NA | endemic | 0 | 1 | 0 | 0 | 0 | 0 | 0 |
| Mantellidae | *Aglyptodactylus* | *Aglyptodactylus inguinalis* | LC | endemic | 0 | 1 | 1 | 0 | 0 | 0 | 0 |
|  |  | *Aglyptodactylus madagascariensis* | LC | endemic | 1 | 1 | 0 | 1 | 0 | 0 | 0 |
|  | *Blommersia* | *Blommersia sp. CaNEW* | NA | endemic | 0 | 1 | 0 | 0 | 0 | 0 | 0 |
|  |  | *Blommersia wittei* | LC | endemic | 0 | 1 | 1 | 1 | 1 | 0 | 0 |
|  | *Boophis* | *Boophis albilabris* | LC | endemic | 0 | 1 | 1 | 1 | 0 | 0 | 0 |
|  |  | *Boophis blommersae* | VU | endemic | 1 | 0 | 0 | 0 | 0 | 0 | 0 |
|  |  | *Boophis englaenderi* | VU | endemic | 1 | 0 | 1 | 0 | 1 | 0 | 0 |
|  |  | *Boophis roseipalmatus* | LC | endemic | 1 | 1 | 0 | 1 | 0 | 0 | 0 |
|  |  | *Boophis septentrionalis* | LC | endemic | 0 | 0 | 0 | 1 | 0 | 0 | 0 |
|  |  | *Boophis sp.* | NA | endemic | 0 | 0 | 1 | 0 | 0 | 0 | 0 |
|  |  | *Boophis sp. CaNEW aff. englaenderi* | NA | endemic | 0 | 0 | 1 | 0 | 0 | 0 | 0 |
|  |  | *Boophis sp. CaNEW aff. rappiodes* | NA | endemic | 0 | 0 | 0 | 1 | 0 | 0 | 0 |
|  |  | *Boophis sp1* | NA | endemic | 0 | 0 | 0 | 1 | 0 | 0 | 0 |
|  |  | *Boophis sp3* | NA | endemic | 0 | 1 | 0 | 1 | 0 | 0 | 0 |
|  |  | *Boophis tephraeomystax* | LC | endemic | 1 | 1 | 1 | 1 | 1 | 1 | 1 |
|  | *Gephyromantis* | *Gephyromantis ambohitra* | VU | endemic | 1 | 0 | 0 | 0 | 0 | 0 | 0 |
|  |  | *Gephyromantis granulatus* | LC | endemic | 1 | 1 | 1 | 0 | 0 | 0 | 0 |
|  |  | *Gephyromantis luteus* | LC | endemic | 1 | 0 | 0 | 0 | 0 | 0 | 0 |
|  |  | *Gephyromantis pseudoasper* | LC | endemic | 1 | 0 | 0 | 0 | 0 | 0 | 0 |
|  |  | *Gephyromantis redimitus* | LC | endemic | 1 | 1 | 0 | 1 | 0 | 0 | 0 |
|  |  | *Gephyromantis sp.Ca17* | NA | endemic | 1 | 1 | 0 | 0 | 0 | 0 | 0 |
|  |  | *Gephyromantis striatus* | VU | endemic | 1 | 0 | 0 | 0 | 0 | 0 | 0 |
|  |  | *Gephyromantis tandroka* | VU | endemic | 1 | 0 | 0 | 0 | 0 | 0 | 0 |
|  | *Guibemantis* | *Guibemantis sp. Ca16* | NA | endemic | 0 | 0 | 10 | 0 | 0 | 0 | 0 |
|  |  | *Guibemantis sp. CaNEW* | NA | endemic | 1 | 0 | 0 | 0 | 0 | 0 | 0 |
|  |  | *Guibemantis tornieri* | LC | endemic | 1 | 0 | 0 | 0 | 0 | 0 | 0 |
|  | *Mantella* | *Mantella ebenaui* | LC | endemic | 0 | 1 | 0 | 0 | 0 | 0 | 0 |
|  |  | *Mantella manery* | VU | endemic | 1 | 0 | 0 | 0 | 0 | 0 | 0 |
|  |  | *Mantella nigricans* | LC | endemic | 1 | 0 | 0 | 0 | 0 | 0 | 0 |
|  | *Mantidactylus* | *Mantidactylus sp. aff.*  *'betsileanus North Clade'* | NA | endemic | 0 | 1 | 0 | 0 | 0 | 0 | 0 |
|  |  | *Mantidactylus sp. Ca12* | NA | endemic | 1 | 0 | 0 | 0 | 0 | 0 | 0 |
|  |  | *Mantidactylus sp. Ca41* | NA | endemic | 1 | 0 | 0 | 0 | 0 | 0 | 0 |
|  |  | *Mantidactylus sp. CaNEW2* | NA | endemic | 0 | 1 | 0 | 0 | 0 | 0 | 0 |
|  |  | *Mantidactylus sp.* | NA | endemic | 0 | 1 | 0 | 0 | 0 | 0 | 0 |
|  | *Spinomantis* | *Spinomantis aglavei* | LC | endemic | 1 | 0 | 0 | 0 | 0 | 0 | 0 |
|  |  | *Spinomantis fimbriatus* | LC | endemic | 1 | 0 | 0 | 0 | 0 | 0 | 0 |
| Microhylidae | *Cophyla* | *Cophyla fortuna* | NE | endemic | 0 | 1 | 0 | 0 | 0 | 0 | 0 |
|  | *Platypelis* | *Platypelis grandis* | LC | endemic | 1 | 0 | 0 | 0 | 0 | 0 | 0 |
|  |  | *Platypelis tetra* | EN | endemic | 1 | 0 | 0 | 0 | 0 | 0 | 0 |
|  | *Plethodontohyla* | *Plethodontohyla notostica* | LC | endemic | 1 | 1 | 0 | 0 | 0 | 0 | 0 |
|  | *Rhombophryne* | *Rhombophryne sp* | NA | endemic | 1 | 0 | 0 | 0 | 0 | 0 | 0 |
|  |  | *Rhombophryne sp. CaNEW* | NA | endemic | 1 | 0 | 0 | 0 | 0 | 0 | 0 |
|  | *Stumpffia* | *Stumpffia achyllei* | NE | endemic | 1 | 0 | 0 | 0 | 0 | 0 | 0 |
|  |  | *Stumpffia diutissuma* | NE | endemic | 1 | 0 | 0 | 0 | 0 | 0 | 0 |
|  |  | *Stumpffia iharana* | NE | endemic | 0 | 1 | 0 | 0 | 0 | 0 | 0 |
|  |  | *Stumpffia mamitika* | NE | endemic | 1 | 1 | 1 | 1 | 1 | 1 | 0 |
|  |  | *Stumpffia sorata* | NE | endemic | 1 | 1 | 0 | 0 | 0 | 0 | 0 |
|  |  | *Stumpffia sp* | NA | endemic | 1 | 0 | 0 | 0 | 0 | 0 | 0 |
|  |  | *Stumpffia sp. aff. edmondsi* | NA | endemic | 0 | 1 | 0 | 0 | 0 | 0 | 0 |
|  |  | *Stumpffia sp. CaNEW1* | NA | endemic | 0 | 1 | 0 | 0 | 0 | 0 | 0 |
|  |  | *Stumpffia sp. CaNEW2* | NA | endemic | 1 | 0 | 0 | 0 | 0 | 0 | 0 |
|  |  | *Stumpffia sp1* | NA | endemic | 0 | 1 | 0 | 0 | 0 | 0 | 0 |
| Ptychadenidae | *Ptychadena* | *Ptychadena mascareniensis* | LC | non-endemic | 0 | 0 | 1 | 1 | 1 | 1 | 1 |
| **Reptile** | | | | | | | | | | | |
| Boidae | *Acrantophis* | *Acranthophis madagascariensis* | LC | endemic | 0 | 1 | 0 | 0 | 0 | 0 | 1 |
|  | *Sanzinia* | *Sanzinia madagascariensis* | LC | non-endemic | 1 | 0 | 0 | 0 | 0 | 0 | 0 |
| Chamaeleonidae | *Brookesia* | *Brookesia griveaudi* | NT | endemic | 1 | 1 | 1 | 0 | 0 | 0 | 0 |
|  |  | *Brookesia karchei* | EN | endemic | 1 | 1 | 0 | 0 | 0 | 0 | 0 |
|  |  | *Brookesia sp.* | NA | endemic | 1 | 1 | 1 | 0 | 0 | 0 | 0 |
|  | *Calumma* | *Calumma cf. nasutum* | NA | endemic | 0 | 0 | 1 | 0 | 0 | 0 | 0 |
|  |  | *Calumma cf. marojezense* | NA | endemic | 1 | 0 | 0 | 0 | 0 | 0 | 0 |
|  |  | *Calumma cucullatum* | VU | endemic | 1 | 0 | 0 | 0 | 0 | 0 | 0 |
|  |  | *Calumma nasutum* | LC | endemic | 1 | 1 | 1 | 1 | 1 | 1 | 0 |
|  |  | *Calumma sp.* | NA | endemic | 1 | 0 | 1 | 0 | 0 | 0 | 0 |
|  | *Furcifer* | *Furcifer pardalis* | LC | endemic | 1 | 1 | 1 | 1 | 1 | 1 | 1 |
|  |  | *Furcifer timoni* | NT | non-endemic | 1 | 0 | 0 | 0 | 0 | 0 | 0 |
| Gekkonidae | *Blaesodactylus* | *Blaesodactylus antongilensis* | LC | endemic | 1 | 0 | 0 | 0 | 0 | 0 | 0 |
|  | *Ebenavia* | *Ebenavia cf. inuinguis* | NA | NA | 1 | 1 | 1 | 1 | 1 | 0 | 0 |
|  | *Geckolepis* | *Geckolepis cf. maculata* | NA | NA | 1 | 1 | 1 | 1 | 1 | 0 | 0 |
|  | *Hemidactylus* | *Hemidactylus sp.* | NA | NA | 0 | 0 | 1 | 0 | 1 | 1 | 0 |
|  | *Lygodactylus* | *Lygodactylus heterurus* | LC | endemic | 0 | 1 | 1 | 1 | 1 | 0 | 0 |
|  |  | *Lygodactylus sp.* | NA | NA | 1 | 1 | 1 | 1 | 1 | 0 | 0 |
|  | *Paroedura* | *Paroedura gracilis* | LC | endemic | 1 | 1 | 0 | 0 | 0 | 0 | 0 |
|  |  | *Paroedura stumpffi* | LC | non-endemic | 0 | 1 | 1 | 0 | 0 | 0 | 0 |
|  | *Phelsuma* | *Phelsuma abbotti* | LC | non-endemic | 0 | 0 | 1 | 0 | 0 | 0 | 0 |
|  |  | *Phelsuma cf. lineata* | NA | NA | 0 | 0 | 0 | 1 | 1 | 0 | 0 |
|  |  | *Phelsuma dorsivittata* | NT | non-endemic | 0 | 0 | 1 | 0 | 0 | 0 | 0 |
|  |  | *Phelsuma guttata* | LC | endemic | 1 | 1 | 1 | 0 | 0 | 0 | 0 |
|  |  | *Phelsuma laticauda* | LC | non-endemic | 0 | 1 | 1 | 1 | 1 | 1 | 1 |
|  |  | *Phelsuma lineata* | LC | endemic | 1 | 1 | 1 | 1 | 1 | 1 | 0 |
|  |  | *Phelsuma madagascariensis* | LC | endemic | 0 | 1 | 1 | 1 | 1 | 1 | 1 |
|  |  | *Phelsuma masohoala* | CR | endemic | 0 | 0 | 0 | 1 | 0 | 0 | 0 |
|  |  | *Phelsuma quadriocellata* | LC | endemic | 1 | 1 | 0 | 1 | 0 | 0 | 0 |
|  |  | *Phelsuma seippi* | EN | endemic | 1 | 1 | 1 | 0 | 1 | 0 | 0 |
|  |  | *Phelsuma sp.* | NA | NA | 0 | 0 | 1 | 0 | 0 | 0 | 0 |
|  | *Uroplatus* | *Uroplatus fimbriatus* | LC | endemic | 1 | 0 | 0 | 0 | 0 | 0 | 0 |
|  |  | *Uroplatus finaritra* | NE | endemic | 1 | 0 | 0 | 0 | 0 | 0 | 0 |
|  |  | *Uroplatus giganteus* | VU | endemic | 1 | 0 | 0 | 0 | 0 | 0 | 0 |
|  |  | *Uroplatus lineatus* | LC | endemic | 1 | 0 | 0 | 0 | 0 | 0 | 0 |
|  |  | *Uroplatus sikorae* | LC | endemic | 1 | 0 | 0 | 0 | 0 | 0 | 0 |
|  |  | *Uroplatus sp. aff. fimbriatus* | NA | endemic | 1 | 0 | 0 | 0 | 0 | 0 | 0 |
| Gerrhosauridae | *Zonosaurus* | *Zonosaurus madagascariensis* | LC | non-endemic | 0 | 1 | 1 | 1 | 1 | 0 | 1 |
|  |  | *Zonosaurus rufipes* | NT | endemic | 1 | 1 | 1 | 0 | 1 | 0 | 0 |
|  |  | *Zonosaurus subinicolor* | EN | endemic | 0 | 1 | 0 | 0 | 0 | 0 | 0 |
| Lamprophiidae | *Alluaudina* | *Alluaudina bellyi* | LC | endemic | 1 | 0 | 0 | 0 | 0 | 0 | 0 |
|  | *Dromicodryas* | *Dromicodryas quadrilineatus* | LC | endemic | 0 | 1 | 1 | 1 | 1 | 0 | 1 |
|  | *Elapotinus* | *Elapotinus picteti* | LC | endemic | 1 | 0 | 0 | 0 | 0 | 0 | 0 |
|  | *Ithycyphus* | *Ithycyphus miniatus* | LC | non-endemic | 0 | 1 | 0 | 1 | 0 | 0 | 0 |
|  |  | *Ithycyphus perineti* | LC | endemic | 1 | 0 | 1 | 0 | 0 | 0 | 0 |
|  | *Langaha* | *Langaha madagascariensis* | LC | endemic | 0 | 1 | 1 | 0 | 0 | 0 | 0 |
|  | *Leioheterodon* | *Leioheterodon madagascariensis* | LC | non-endemic | 0 | 0 | 1 | 0 | 1 | 0 | 1 |
|  |  | *Leioheterodon modestus* | LC | endemic | 0 | 0 | 0 | 1 | 0 | 0 | 1 |
|  | *Liophidium* | *Liophidium torquatum* | LC | endemic | 1 | 0 | 0 | 1 | 0 | 0 | 0 |
|  | *Lycodryas* | *Lycodryas granuliceps* | LC | endemic | 0 | 0 | 1 | 0 | 1 | 0 | 0 |
|  | *Madagascarophis* | *Madagascarophis colubrinus* | LC | endemic | 0 | 0 | 0 | 1 | 0 | 0 | 0 |
|  | *Pseudoxyrhopus* | *Pseudoxyrhopus cf. sokosoko* | NA | endemic | 0 | 1 | 0 | 0 | 0 | 0 | 0 |
|  |  | *Pseudoxyrhopus heterurus* | LC | endemic | 1 | 0 | 0 | 0 | 0 | 0 | 0 |
|  |  | *Pseudoxyrhopus sp.* | NA | endemic | 0 | 1 | 0 | 0 | 0 | 0 | 0 |
|  | *Thamnosophis* | *Thamnosophis lateralis* | LC | endemic | 0 | 1 | 1 | 1 | 0 | 0 | 1 |
|  |  | *Thamnosophis martae* | EN | endemic | 0 | 0 | 0 | 1 | 0 | 0 | 0 |
|  |  | *Thamnosophis sp.* | NA | endemic | 1 | 0 | 0 | 0 | 0 | 0 | 0 |
| Scincidae | *Amphiglossus* | *Amphiglossus cf spilostichus* | NA | NA | 1 | 0 | 0 | 0 | 0 | 0 | 0 |
|  | *Madascincus* | *Madascincus minutus* | LC | endemic | 1 | 1 | 0 | 0 | 0 | 0 | 0 |
|  | *Trachylepis* | *Trachylepis elegans* | LC | endemic | 0 | 1 | 1 | 0 | 1 | 1 | 1 |
|  |  | *Trachylepis gravenhorstii* | LC | endemic | 0 | 1 | 1 | 1 | 1 | 1 | 1 |

**Table SI 2**: Tukey post-hoc pairwise comparisons for amphibians and reptiles between land-use types. **OGF** (Old-growth forest), **FF** (forest fragment), **VFST** (Forest-derived vanilla agroforest), **VFLW** (Fallow-derived vanilla agroforest), **WF** (woody fallow), **HF** (herbaceous fallow), **RP** (rice paddy). Significant p-values (<0.05) are highlighted in bold.

| Comparison | Estimate | Std. Error | z value | Pr(>\|z\|) | sign. |
| --- | --- | --- | --- | --- | --- |
| **Amphibian** | | | | | |
| FF - OGF | -0.601 | 0.174 | -3.448 | **<0.001** | *** |
| VFST - OGF | -0.704 | 0.180 | -3.905 | **<0.001** | *** |
| VFLW - OGF | -0.715 | 0.147 | -4.848 | **<0.001** | *** |
| WF - OGF | -0.869 | 0.190 | -4.555 | **<0.001** | *** |
| HF - OGF | -1.099 | 0.207 | -5.297 | **<0.001** | *** |
| RP - OGF | -1.588 | 0.252 | -6.308 | **<0.001** | *** |
| VFST - FF | -0.103 | 0.203 | -0.507 | 0.611 |  |
| VFLW - FF | -0.114 | 0.175 | -0.652 | 0.514 |  |
| WF - FF | -0.268 | 0.213 | -1.261 | 0.207 |  |
| HF - FF | -0.498 | 0.228 | -2.186 | **0.028** | * |
| RP - FF | -0.987 | 0.269 | -3.674 | **<0.001** | *** |
| VFLW - VFST | -0.011 | 0.181 | -0.060 | 0.952 |  |
| WF - VFST | -0.165 | 0.218 | -0.758 | 0.448 |  |
| HF - VFST | -0.395 | 0.232 | -1.698 | 0.089 | . |
| RP - VFST | -0.884 | 0.273 | -3.242 | **0.001** | ** |
| WF - VFLW | -0.154 | 0.191 | -0.805 | 0.420 |  |
| HF - VFLW | -0.384 | 0.208 | -1.845 | 0.065 | . |
| RP - VFLW | -0.873 | 0.252 | -3.462 | **<0.001** | *** |
| HF - WF | -0.230 | 0.241 | -0.954 | 0.340 |  |
| RP - WF | -0.719 | 0.280 | -2.570 | **0.010** | * |
| RP - HF | -0.490 | 0.291 | -1.680 | 0.093 | . |
| **Reptile** | | | | | |
| FF - OGF | -0.021 | 0.144 | -0.144 | 0.886 |  |
| VFST - OGF | -0.041 | 0.144 | -0.289 | 0.773 |  |
| VFLW - OGF | -0.358 | 0.132 | -2.707 | **0.007** | ** |
| WF - OGF | -0.426 | 0.161 | -2.651 | **0.008** | ** |
| HF - OGF | -1.030 | 0.197 | -5.229 | **<0.001** | *** |
| RP - OGF | -1.366 | 0.224 | -6.097 | **<0.001** | *** |
| VFST - FF | -0.021 | 0.145 | -0.145 | 0.885 |  |
| VFLW - FF | -0.338 | 0.133 | -2.536 | **0.011** | * |
| WF - FF | -0.405 | 0.161 | -2.513 | **0.012** | * |
| HF - FF | -1.009 | 0.197 | -5.110 | **<0.001** | *** |
| RP - FF | -1.345 | 0.225 | -5.992 | **<0.001** | *** |
| VFLW - VFST | -0.316 | 0.134 | -2.363 | **0.018** | * |
| WF - VFST | -0.384 | 0.162 | -2.372 | **0.018** | * |
| HF - VFST | -0.988 | 0.198 | -4.989 | **<0.001** | *** |
| RP - VFST | -1.324 | 0.225 | -5.886 | **<0.001** | *** |
| WF - VFLW | -0.068 | 0.151 | -0.449 | 0.654 |  |
| HF - VFLW | -0.671 | 0.189 | -3.545 | **<0.001** | *** |
| RP - VFLW | -1.008 | 0.217 | -4.635 | **<0.001** | *** |
| HF - WF | -0.604 | 0.210 | -2.871 | **0.004** | ** |
| RP - WF | -0.940 | 0.236 | -3.986 | **<0.001** | *** |
| RP - HF | -0.336 | 0.262 | -1.285 | 0.199 |  |
| Signif. codes: 0 ‘***’ 0.001 ‘**’ 0.01 ‘*’ 0.05 ‘.’ 0.1 ‘ ’ 1 | | | | | |

**Table SI 3**: Extrapolated species diversity (species richness, Shannon diversity and Simpson diversity) based on 5000 encounters and includes the lower and upper 95% confidence interval for amphibians and reptiles. **m**: sample size, **SC**: sample coverage estimate, **qD**: diversity estimate of order q, qD.LCL: the 95% lower confidence limits of diversity, qD.UCL: the 95% upper confidence limits of diversity. **OGF** (Old-growth forest), **FF** (forest fragment), **VFST** (Forest-derived vanilla agroforest), **VFLW** (Fallow-derived vanilla agroforest), **WF** (woody fallow), **HF** (herbaceous fallow), **RP** (rice paddy).

| LUT | m | method | order | SC | qD | qD.LCL | qD.UCL |
| --- | --- | --- | --- | --- | --- | --- | --- |
| **Amphibian** | | | | | | | |
| OGF | 5000 | extrapolated | 0 | 1 | 59.971 | 20.356 | 99.585 |
| OGF | 5000 | extrapolated | 1 | 1 | 11.615 | 10.009 | 13.221 |
| OGF | 5000 | extrapolated | 2 | 1 | 6.869 | 6.024 | 7.715 |
| FF | 5000 | extrapolated | 0 | 1 | 46.108 | 12.410 | 79.807 |
| FF | 5000 | extrapolated | 1 | 1 | 6.542 | 5.451 | 7.633 |
| FF | 5000 | extrapolated | 2 | 1 | 3.639 | 3.232 | 4.045 |
| VFST | 5000 | extrapolated | 0 | 1 | 17.986 | 9.082 | 26.890 |
| VFST | 5000 | extrapolated | 1 | 1 | 6.636 | 5.919 | 7.353 |
| VFST | 5000 | extrapolated | 2 | 1 | 4.939 | 4.350 | 5.528 |
| VFLW | 5000 | extrapolated | 0 | 1 | 24.970 | 7.005 | 42.935 |
| VFLW | 5000 | extrapolated | 1 | 1 | 5.552 | 4.966 | 6.139 |
| VFLW | 5000 | extrapolated | 2 | 1 | 4.072 | 3.680 | 4.464 |
| WF | 5000 | extrapolated | 0 | 1 | 8.996 | 7.375 | 10.616 |
| WF | 5000 | extrapolated | 1 | 1 | 4.843 | 4.446 | 5.240 |
| WF | 5000 | extrapolated | 2 | 1 | 4.135 | 3.707 | 4.562 |
| HF | 5000 | extrapolated | 0 | 1 | 6.498 | 3.898 | 9.099 |
| HF | 5000 | extrapolated | 1 | 1 | 3.492 | 3.212 | 3.771 |
| HF | 5000 | extrapolated | 2 | 1 | 3.128 | 2.878 | 3.379 |
| RP | 5000 | extrapolated | 0 | 1 | 4.998 | 4.348 | 5.648 |
| RP | 5000 | extrapolated | 1 | 1 | 1.331 | 1.276 | 1.386 |
| RP | 5000 | extrapolated | 2 | 1 | 1.169 | 1.130 | 1.208 |
| **Reptile** | | | | | | | |
| OGF | 5000 | extrapolated | 0 | 1 | 41.963 | 20.405 | 63.522 |
| OGF | 5000 | extrapolated | 1 | 1 | 16.282 | 13.376 | 19.188 |
| OGF | 5000 | extrapolated | 2 | 1 | 8.267 | 6.192 | 10.341 |
| FF | 5000 | extrapolated | 0 | 1 | 40.048 | 10.192 | 69.904 |
| FF | 5000 | extrapolated | 1 | 1 | 15.463 | 13.557 | 17.368 |
| FF | 5000 | extrapolated | 2 | 1 | 10.935 | 9.614 | 12.256 |
| VFST | 5000 | extrapolated | 0 | 1 | 35.318 | 18.210 | 52.426 |
| VFST | 5000 | extrapolated | 1 | 1 | 14.355 | 12.633 | 16.077 |
| VFST | 5000 | extrapolated | 2 | 1 | 9.323 | 7.998 | 10.647 |
| VFLW | 5000 | extrapolated | 0 | 1 | 26.949 | 8.198 | 45.699 |
| VFLW | 5000 | extrapolated | 1 | 1 | 6.902 | 6.268 | 7.535 |
| VFLW | 5000 | extrapolated | 2 | 1 | 5.075 | 4.631 | 5.519 |
| WF | 5000 | extrapolated | 0 | 1 | 46.603 | 11.526 | 81.679 |
| WF | 5000 | extrapolated | 1 | 1 | 8.715 | 7.464 | 9.965 |
| WF | 5000 | extrapolated | 2 | 1 | 6.236 | 5.415 | 7.057 |
| HF | 5000 | extrapolated | 0 | 1 | 8.000 | 6.830 | 9.170 |
| HF | 5000 | extrapolated | 1 | 1 | 3.227 | 2.748 | 3.705 |
| HF | 5000 | extrapolated | 2 | 1 | 2.194 | 1.880 | 2.507 |
| RP | 5000 | extrapolated | 0 | 1 | 18.862 | 4.312 | 33.413 |
| RP | 5000 | extrapolated | 1 | 1 | 6.046 | 3.255 | 8.836 |
| RP | 5000 | extrapolated | 2 | 1 | 3.129 | 1.790 | 4.468 |

**Table SI 4**: Pairwise-adonis for amphibian and reptile species composition between land-use types. **OGF** (Old-growth forest), **FF** (forest fragment), **VFST** (Forest-derived vanilla agroforest), **VFLW** (Fallow-derived vanilla agroforest), **WF** (woody fallow), **HF** (herbaceous fallow), **RP** (rice paddy). Significant p-values (<0.05) are highlighted in bold.

| **number** | **pairs** | **Df** | **SumsOfSqs** | **F.Model** | **R2** | **p.value** | **p.adjusted** | **Sign.** |
| --- | --- | --- | --- | --- | --- | --- | --- | --- |
| **Amphibian** | | | | | | | | |
| 1 | OGF vs FF | 1 | 3.176 | 11.689 | 0.394 | 0.001 | **0.021** | . |
| 2 | OGF vs HF | 1 | 5.913 | 27.739 | 0.620 | 0.001 | **0.021** | . |
| 3 | OGF vs RP | 1 | 7.904 | 54.608 | 0.752 | 0.001 | **0.021** | . |
| 4 | OGF vs VFST | 1 | 4.645 | 17.771 | 0.497 | 0.001 | **0.021** | . |
| 5 | OGF vs VFLW | 1 | 6.789 | 32.051 | 0.534 | 0.001 | **0.021** | . |
| 6 | OGF vs WF | 1 | 5.297 | 23.929 | 0.571 | 0.001 | **0.021** | . |
| 7 | FF vs HF | 1 | 1.534 | 6.723 | 0.283 | 0.001 | **0.021** | . |
| 8 | FF vs RP | 1 | 2.956 | 18.595 | 0.508 | 0.001 | **0.021** | . |
| 9 | FF vs VFST | 1 | 0.547 | 1.986 | 0.099 | 0.037 | 0.777 |  |
| 10 | FF vs VFLW | 1 | 1.322 | 5.983 | 0.176 | 0.001 | **0.021** | . |
| 11 | FF vs WF | 1 | 0.989 | 4.198 | 0.189 | 0.001 | **0.021** | . |
| 12 | HF vs RP | 1 | 1.418 | 15.112 | 0.471 | 0.001 | **0.021** | . |
| 13 | HF vs VFST | 1 | 0.678 | 3.118 | 0.155 | 0.008 | 0.168 |  |
| 14 | HF vs VFLW | 1 | 0.282 | 1.548 | 0.054 | 0.147 | 1 |  |
| 15 | HF vs WF | 1 | 0.290 | 1.657 | 0.089 | 0.153 | 1 |  |
| 16 | RP vs VFST | 1 | 2.508 | 16.868 | 0.484 | 0.001 | **0.021** | . |
| 17 | RP vs VFLW | 1 | 2.932 | 21.037 | 0.429 | 0.001 | **0.021** | . |
| 18 | RP vs WF | 1 | 2.162 | 19.909 | 0.525 | 0.001 | **0.021** | . |
| 19 | VFST vs VFLW | 1 | 0.244 | 1.138 | 0.039 | 0.335 | 1 |  |
| 20 | VFST vs WF | 1 | 0.259 | 1.147 | 0.060 | 0.330 | 1 |  |
| 21 | VFLW vs WF | 1 | 0.107 | 0.568 | 0.020 | 0.823 | 1 |  |
| **Reptile** | | | | | | | | |
| 1 | OGF vs FF | 1 | 1.038 | 4.269 | 0.192 | 0.001 | **0.021** | . |
| 2 | OGF vs HF | 1 | 5.596 | 29.803 | 0.623 | 0.001 | **0.021** | . |
| 3 | OGF vs RP | 1 | 6.956 | 20.906 | 0.537 | 0.001 | **0.021** | . |
| 4 | OGF vs VFST | 1 | 1.860 | 7.210 | 0.286 | 0.001 | **0.021** | . |
| 5 | OGF vs VFLW | 1 | 4.763 | 23.014 | 0.451 | 0.001 | **0.021** | . |
| 6 | OGF vs WF | 1 | 3.462 | 15.205 | 0.458 | 0.001 | **0.021** | . |
| 7 | FF vs HF | 1 | 2.392 | 13.454 | 0.428 | 0.001 | **0.021** | . |
| 8 | FF vs RP | 1 | 2.042 | 6.328 | 0.260 | 0.001 | **0.021** | . |
| 9 | FF vs VFST | 1 | 0.600 | 2.420 | 0.119 | 0.002 | **0.042** | . |
| 10 | FF vs VFLW | 1 | 1.704 | 8.495 | 0.233 | 0.001 | **0.021** | . |
| 11 | FF vs WF | 1 | 1.243 | 5.707 | 0.241 | 0.001 | **0.021** | . |
| 12 | HF vs RP | 1 | 0.476 | 1.782 | 0.090 | 0.080 | 1 |  |
| 13 | HF vs VFST | 1 | 1.681 | 8.731 | 0.327 | 0.001 | **0.021** | . |
| 14 | HF vs VFLW | 1 | 1.291 | 7.830 | 0.219 | 0.001 | **0.021** | . |
| 15 | HF vs WF | 1 | 0.779 | 4.799 | 0.210 | 0.002 | **0.042** | . |
| 16 | RP vs VFST | 1 | 1.380 | 4.089 | 0.185 | 0.001 | **0.021** | . |
| 17 | RP vs VFLW | 1 | 0.935 | 3.623 | 0.115 | 0.002 | **0.042** | . |
| 18 | RP vs WF | 1 | 1.044 | 3.397 | 0.159 | 0.011 | 0.231 |  |
| 19 | VFST vs VFLW | 1 | 0.782 | 3.724 | 0.117 | 0.002 | **0.042** | . |
| 20 | VFST vs WF | 1 | 0.504 | 2.168 | 0.107 | 0.016 | 0.336 |  |
| 21 | VFLW vs WF | 1 | 0.159 | 0.837 | 0.029 | 0.606 | 1 |  |

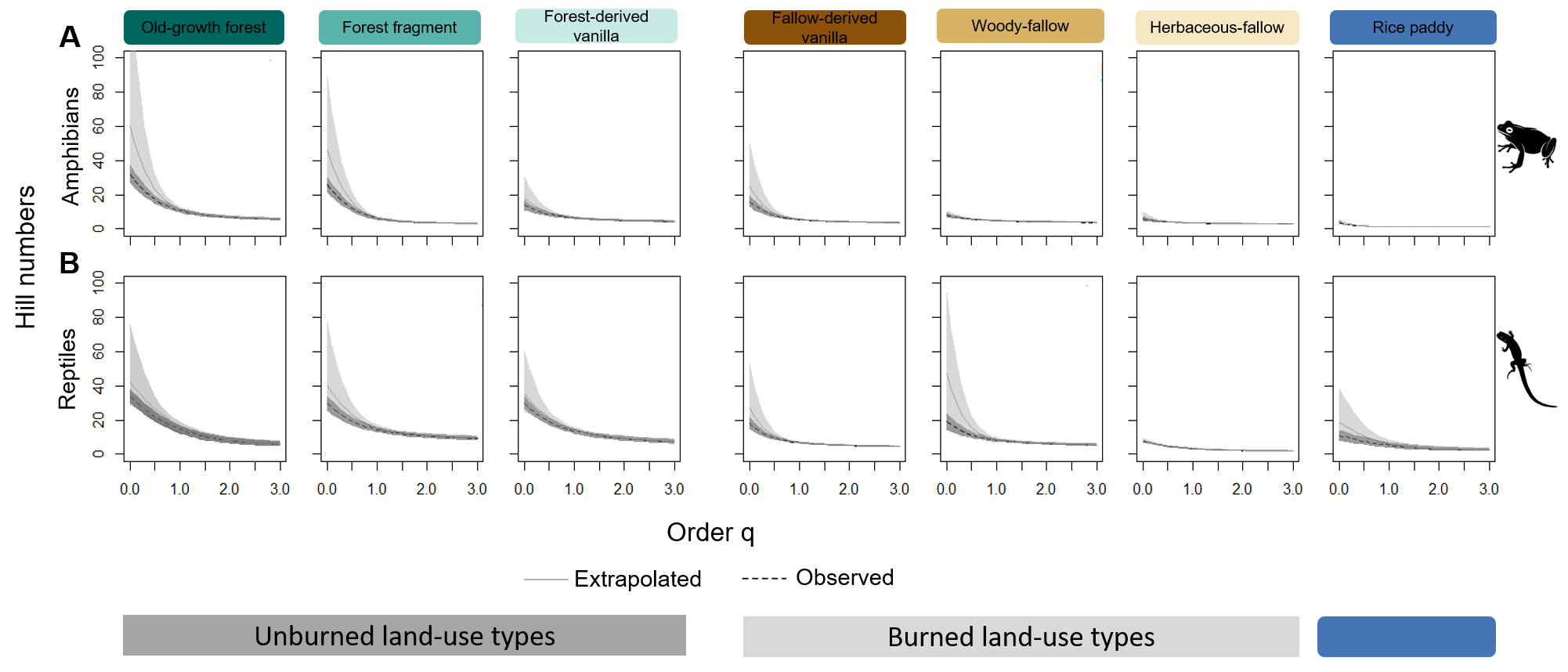

**Figure SI 5**: Curves showing the evenness of (A) amphibian and (B) reptile across seven land-use types in north-eastern Madagascar for orders of q between 0 and 3 with 95% confidence intervals. The steeper the drop is across orders of q, the more uneven a community is. The dotted lines represent the observed diversity and solid lines represent extrapolated diversity. Icon source: Free Icon Library (see reference below).

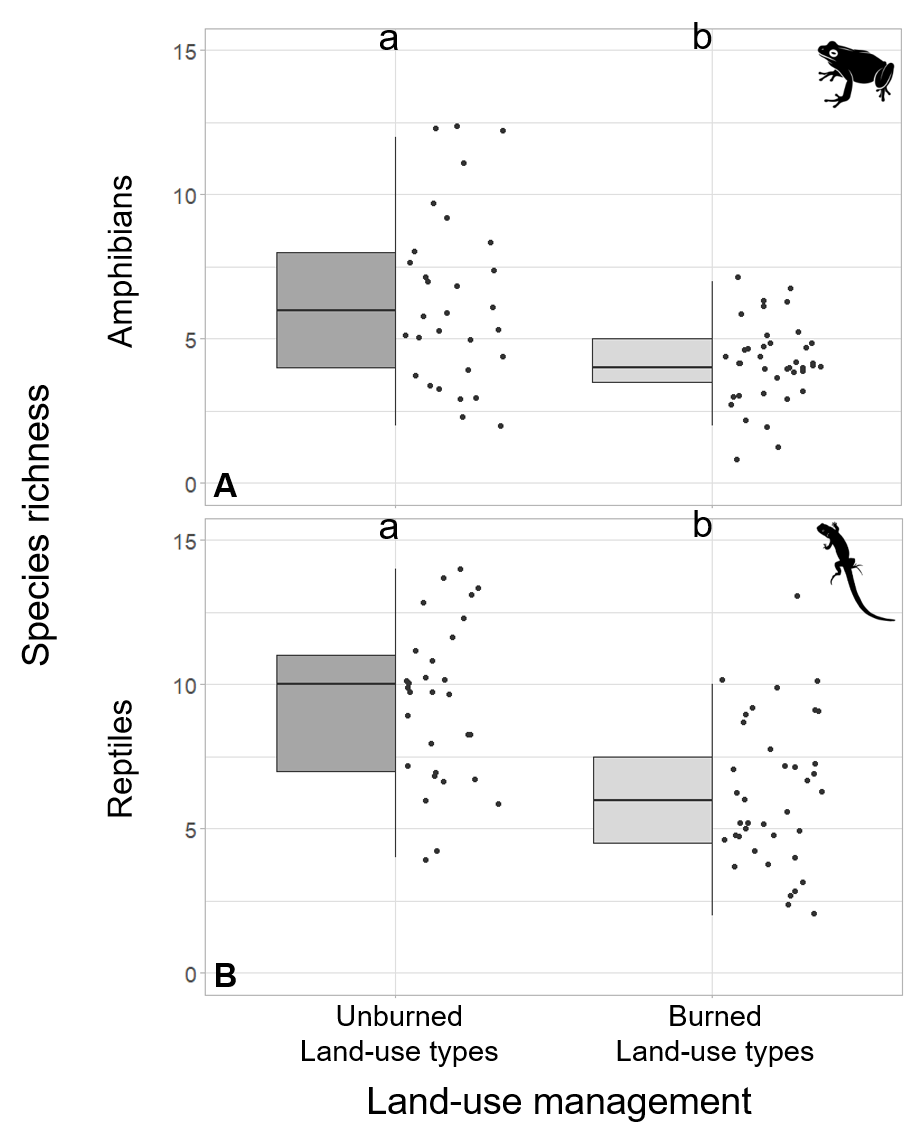

**Figure SI 6:** Plot-level observed amphibian (A) and reptile (B) species richness of unburned land-use types (old-growth forest, forest fragment, and forest-derived vanilla agroforest) and burned land-use types (herbaceous fallow, woody fallow, and fallow-derived vanilla agroforest) in north-eastern Madagascar. The dots represent the species richness per plot in each land-use type. The black horizontal line in the box shows the median. Land-use types with different letters differ significantly based on the Tukey HSD approach (Numeric results in SI: amphibians table SI 7 & reptiles table SI 8). Icon source: Free Icon Library (see reference below).

**Table SI 7**: Comparison of amphibian species richness between unburned and burned land-use types. Significant p-values (<0.05) are highlighted in bold.

|  | Df | Sum Sq | Mean Sq | F value | Pr(>F) | Sign. |
| --- | --- | --- | --- | --- | --- | --- |
| Land-use management | 1 | 78.1 | 78.1 | 15.89 | **<0.001** | *** |
| Residuals | 67 | 329.4 | 4.92 |  |  |  |

Signif. codes: 0 ‘***’ 0.001 ‘**’ 0.01 ‘*’ 0.05 ‘.’ 0.1 ‘ ’ 1

**Table SI 8**. Comparison of reptile species richness between unburned and burned land-use types. Significant p-values (<0.05) are highlighted in bold.

|  | Df | Sum Sq | Mean Sq | F value | Pr(>F) | Sign. |
| --- | --- | --- | --- | --- | --- | --- |
| Land-use management | 1 | 175 | 175.03 | 25.6 | **<0.001** | *** |
| Residuals | 67 | 458 | 6.84 |  |  |  |

Signif. codes: 0 ‘***’ 0.001 ‘**’ 0.01 ‘*’ 0.05 ‘.’ 0.1 ‘ ’ 1

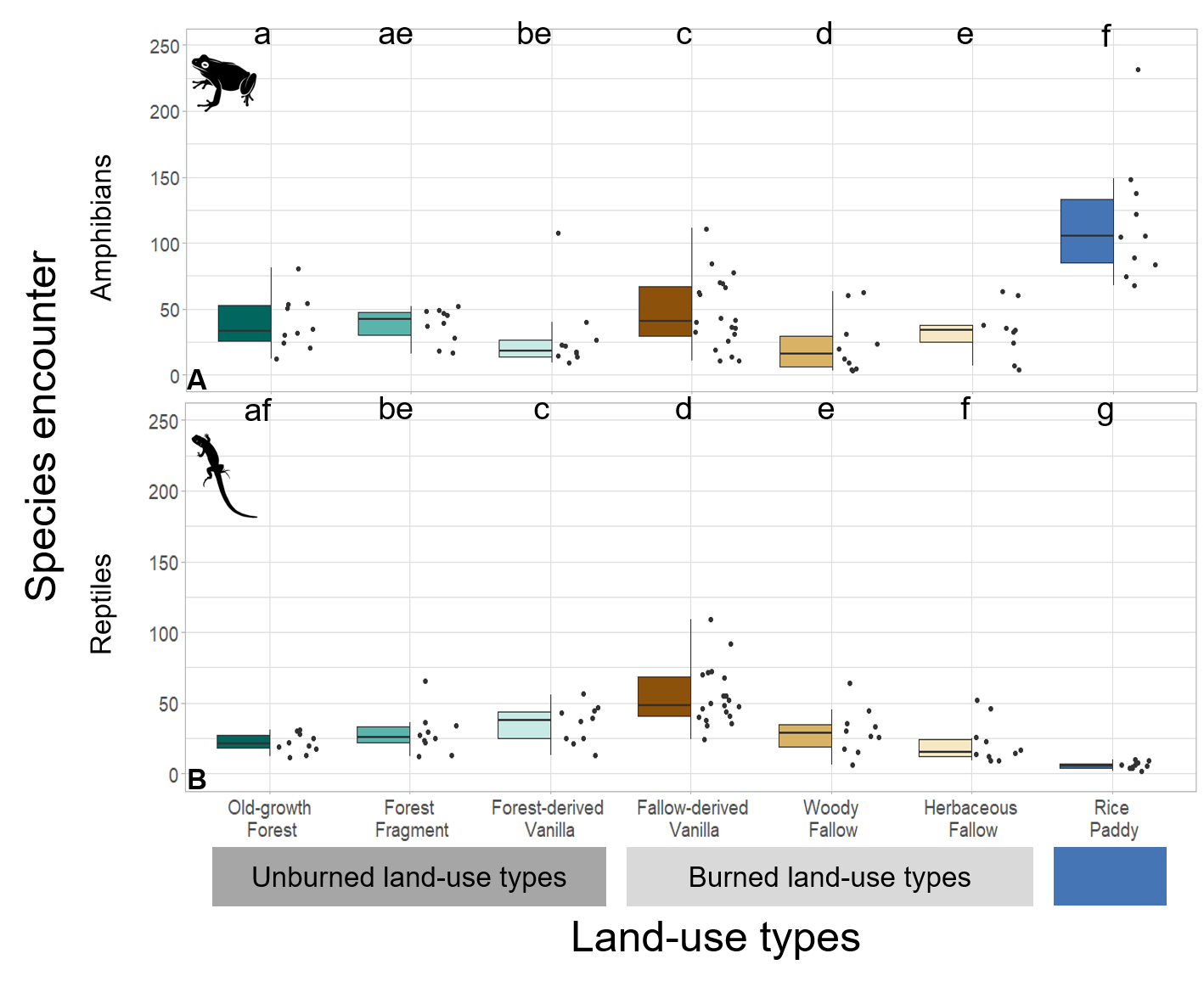

**Figure SI 9:** Plot-level amphibian (A) and reptile (B) species encounters across seven land-use types in north-eastern Madagascar. The dots represent the species encounters per plot in each land-use type. The black horizontal line in the box shows the median. Land-use types with letters in common did not differ significantly based on pairwise comparisons that controlled for inflated false positive errors using the Tukey HSD approach (Numeric results in SI: amphibians & reptiles SI 12). Icon source: Free Icon Library (see reference below).

**Table SI 10:** Comparison of amphibian species encounters across seven land-use types. Significant p-values (<0.05) are highlighted in bold.

|  | Df | Sum Sq | Mean Sq | F value | Pr(>F) | Sign. |
| --- | --- | --- | --- | --- | --- | --- |
| Land-use types | 6 | 60488 | 10081 | 13.01 | **<0.001** | *** |
| Residuals | 72 | 55800 | 775 |  |  |  |

Signif. codes: 0 ‘***’ 0.001 ‘**’ 0.01 ‘*’ 0.05 ‘.’ 0.1 ‘ ’ 1

**Table SI 11:** Comparison of reptile species encounters across seven land-use types. Significant p-values (<0.05) are highlighted in bold.

|  | Df | Sum Sq | Mean Sq | F value | Pr(>F) | Sign. |
| --- | --- | --- | --- | --- | --- | --- |
| Land-use types | 6 | 19311 | 3218 | 14 | **<0.001** | *** |
| Residuals | 73 | 16781 | 230 |  |  |  |

Signif. codes: 0 ‘***’ 0.001 ‘**’ 0.01 ‘*’ 0.05 ‘.’ 0.1 ‘ ’ 1

**Table SI 12:** Tukey post-hoc pairwise comparisons for amphibian and reptile species encountered between land-use types. **OGF** (Old-growth forest), **FF** (forest fragment), **VFST** (Forest-derived vanilla agroforest), **VFLW** (Fallow-derived vanilla agroforest), **WF** (woody fallow), **HF** (herbaceous fallow), **RP** (rice paddy). Significant p-values (<0.05) are highlighted in bold.

| Comparison | Estimate | Std. Error | z value | Pr(>\|z\|) | Sign. |
| --- | --- | --- | --- | --- | --- |
| **Amphibian** | | | | | |
| FF vs OGF | -0.036 | 0.072 | -0.504 | 0.614 |  |
| VFST vs OGF | -0.314 | 0.078 | -4.048 | **<0.001** | *** |
| VFLW vs OGF | 0.179 | 0.060 | 2.979 | **<0.001** | ** |
| WF vs OGF | -0.531 | 0.083 | -6.410 | **<0.001** | *** |
| HF vs OGF | -0.168 | 0.077 | -2.189 | **0.029** | * |
| RP vs OGF | 1.087 | 0.058 | 18.628 | **<0.001** | *** |
| VFST vs FF | -0.278 | 0.078 | -3.553 | **<0.001** | *** |
| VFLW vs FF | 0.215 | 0.061 | 3.537 | **<0.001** | *** |
| WF vs FF | -0.495 | 0.083 | -5.932 | **<0.001** | *** |
| HF vs FF | -0.132 | 0.077 | -1.703 | 0.089 | . |
| RP vs FF | 1.123 | 0.059 | 18.990 | **<0.001** | *** |
| VFLW vs VFST | 0.493 | 0.067 | 7.314 | **<0.001** | *** |
| WF vs VFST | -0.217 | 0.088 | -2.456 | **0.014** | * |
| HF vs VFST | 0.146 | 0.083 | 1.771 | 0.077 | . |
| RP vs VFST | 1.401 | 0.066 | 21.260 | **<0.001** | *** |
| WF vs VFLW | -0.710 | 0.073 | -9.673 | **<0.001** | *** |
| HF vs VFLW | -0.347 | 0.066 | -5.225 | **<0.001** | *** |
| RP vs VFLW | 0.908 | 0.044 | 20.704 | **<0.001** | *** |
| HF vs WF | 0.363 | 0.088 | 4.148 | **<0.001** | *** |
| RP vs WF | 1.618 | 0.072 | 22.466 | **<0.001** | *** |
| RP vs HF | 1.255 | 0.065 | 19.353 | **<0.001** | *** |
| **Reptile** | | | | | |
| FF vs OGF | 0.278 | 0.090 | 3.102 | **0.002** | ** |
| VFST vs OGF | 0.471 | 0.086 | 5.451 | **<0.001** | *** |
| VFLW vs OGF | 0.917 | 0.074 | 12.363 | **<0.001** | *** |
| WF vs OGF | 0.313 | 0.089 | 3.508 | **<0.001** | *** |
| HF vs OGF | 0.005 | 0.096 | 0.048 | 0.962 |  |
| RP vs OGF | -1.324 | 0.148 | -8.962 | **<0.001** | *** |
| VFST vs FF | 0.192 | 0.080 | 2.413 | 0.016 | * |
| VFLW vs FF | 0.639 | 0.066 | 9.642 | **<0.001** | *** |
| WF vs FF | 0.034 | 0.083 | 0.413 | 0.680 |  |
| HF vs FF | -0.274 | 0.090 | -3.055 | 0.002 | ** |
| RP vs FF | -1.603 | 0.144 | -11.135 | **<0.001** | *** |
| VFLW vs VFST | 0.447 | 0.061 | 7.263 | **<0.001** | *** |
| WF vs VFST | -0.158 | 0.079 | -2.003 | 0.045 | * |
| HF vs VFST | -0.466 | 0.086 | -5.406 | **<0.001** | *** |
| RP vs VFST | -1.795 | 0.142 | -12.656 | **<0.001** | *** |
| WF vs VFLW | -0.605 | 0.065 | -9.250 | **<0.001** | *** |
| HF vs VFLW | -0.913 | 0.074 | -12.325 | **<0.001** | *** |
| RP vs VFLW | -2.241 | 0.135 | -16.633 | **<0.001** | *** |
| HF vs WF | -0.308 | 0.089 | -3.461 | **<0.001** | *** |
| RP vs WF | -1.637 | 0.144 | -11.404 | **<0.001** | *** |
| RP vs HF | -1.329 | 0.148 | -8.997 | **<0.001** | *** |

Signif. codes: 0 ‘***’ 0.001 ‘**’ 0.01 ‘*’ 0.05 ‘.’ 0.1 ‘ ’ 1

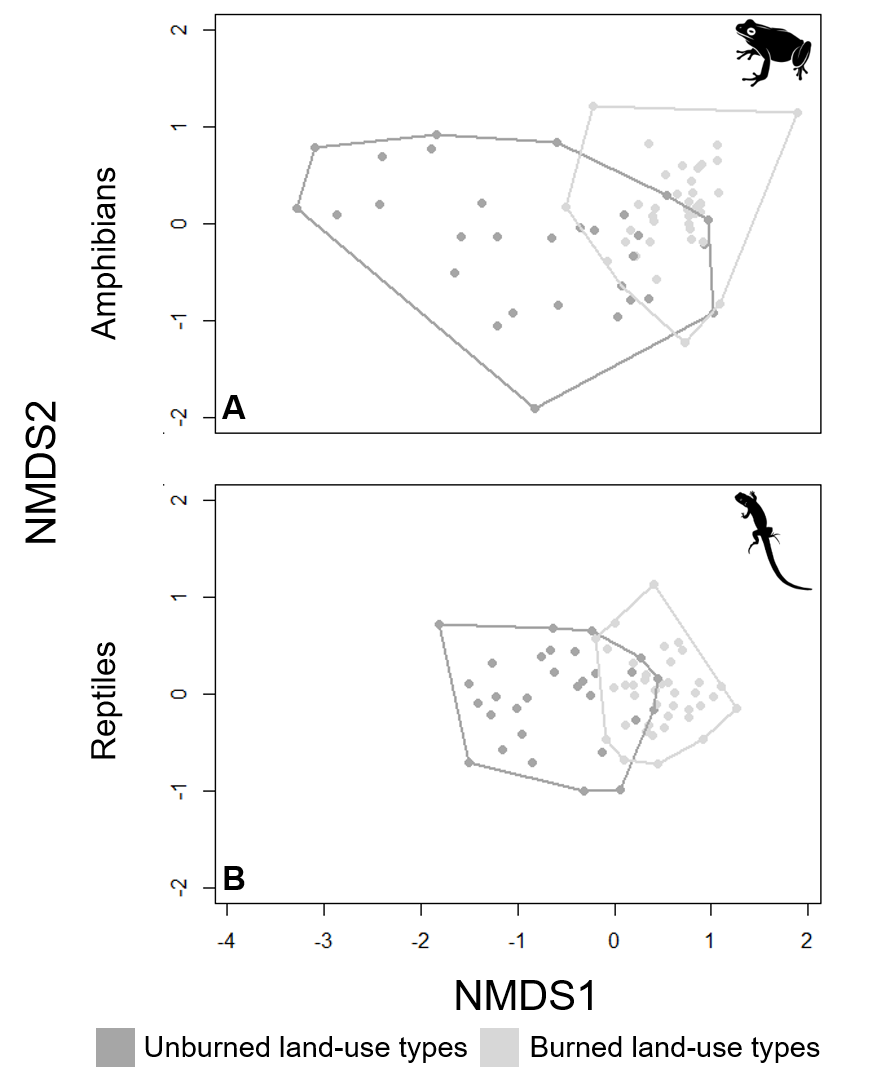

**Figure SI 13:** Species composition in unburned and burned land-use in north-eastern Madagascar. Non-metric dimensional scaling (NMDS) showing community dissimilarity of (A) amphibian and (B) reptile communities. Icon source: Free Icon Library (see reference below).

**Table SI** **14:** Adonis test of community dissimilarity for amphibian species between unburned and burned land-use types. Significant p-values (<0.05) are highlighted in bold.

|  | Df | SumsOfSqs | MeanSqs | F.Model | R2 | Pr(>F) | Sign. |
| --- | --- | --- | --- | --- | --- | --- | --- |
| Land-use management | 1 | 5.524 | 5.524 | 18.389 | 0.193 | **0.001** | *** |
| Residuals | 77 | 23.132 | 0.300 |  | 0.807 |  |  |
| Total | 78 | 28.657 |  |  | 1 |  |  |

Signif. codes: 0 ‘***’ 0.001 ‘**’ 0.01 ‘*’ 0.05 ‘.’ 0.1 ‘ ’ 1

**Table SI** **15:** Adonis test of community dissimilarity for reptile species between unburned and burned land-use types. Significant p-values (<0.05) are highlighted in bold.

|  | Df | SumsOfSqs | MeanSqs | F.Model | R2 | Pr(>F) | Sign |
| --- | --- | --- | --- | --- | --- | --- | --- |
| Land-use management | 1 | 5.946 | 5.946 | 20.777 | 0.210 | **0.001** | *** |
| Residuals | 78 | 22.322 | 0.286 |  | 0.790 |  |  |
| Total | 79 | 28.268 |  |  | 1 |  |  |

Signif. codes: 0 ‘***’ 0.001 ‘**’ 0.01 ‘*’ 0.05 ‘.’ 0.1 ‘ ’ 1

**Table SI 16:** Permutation test for homogeneity of multivariate dispersions for amphibian species composition within unburned and burned land-use types. Significant p-values (<0.05) are highlighted in bold.

|  | Df | Sum Sq | Mean Sq | F | N.Perm | Pr(>F) | Sin. |
| --- | --- | --- | --- | --- | --- | --- | --- |
| Land-use management | 1 | 0.642 | 0.642 | 34.718 | 999 | **0.001** | *** |
| Residuals | 77 | 1.424 | 0.019 |  |  |  |  |

Signif. codes: 0 ‘***’ 0.001 ‘**’ 0.01 ‘*’ 0.05 ‘.’ 0.1 ‘ ’ 1

**Table SI 17:** Permutation test for homogeneity of multivariate dispersions for reptile species composition within unburned and burned land-use types.

|  | Df | Sum Sq | Mean Sq | F | N.Perm | Pr(>F) | Sign. |
| --- | --- | --- | --- | --- | --- | --- | --- |
| Land-use management | 1 | 0.032 | 0.032 | 3.918 | 999 | 0.057 | . |
| Residuals | 78 | 0.640 | 0.008 |  |  |  |  |

Signif. codes: 0 ‘***’ 0.001 ‘**’ 0.01 ‘*’ 0.05 ‘.’ 0.1 ‘ ’ 1

**Table SI 18:** Permutation test for homogeneity of multivariate dispersions for amphibian species composition within seven land-use types. Significant p-values (<0.05) are highlighted in bold.

|  | Df | Sum Sq | Mean Sq | F | N.Perm | Pr(>F) |  |
| --- | --- | --- | --- | --- | --- | --- | --- |
| Land-use types | 6 | 0.519 | 0.087 | 4.400 | 999 | **0.004** | ** |
| Residuals | 72 | 1.417 | 0.020 |  |  |  |  |

Signif. codes: 0 ‘***’ 0.001 ‘**’ 0.01 ‘*’ 0.05 ‘.’ 0.1 ‘ ’ 1

**Table SI 19:** Tukey multiple comparisons of means for homogeneity of multivariate dispersions for amphibian species composition within seven land-use types. **OGF** (Old-growth forest), **FF** (forest fragment), **VFST** (Forest-derived vanilla agroforest), **VFLW** (Fallow-derived vanilla agroforest), **WF** (woody fallow), **HF** (herbaceous fallow), **RP** (rice paddy). Significant p-values (<0.05) are highlighted in bold.

| pairs | difference | lower | upper | p adjusted |
| --- | --- | --- | --- | --- |
| HF vs FF | -0.208 | -0.404 | -0.013 | **0.029** |
| OGF vs FF | -0.005 | -0.195 | 0.185 | 1.000 |
| RP vs FF | -0.224 | -0.415 | -0.034 | **0.011** |
| VFLW vs FF | -0.134 | -0.299 | 0.030 | 0.184 |
| VFST vs FF | -0.035 | -0.225 | 0.156 | 0.998 |
| WF vs FF | -0.083 | -0.273 | 0.107 | 0.837 |
| OGF vs HF | 0.203 | 0.008 | 0.399 | **0.036** |
| RP vs HF | -0.016 | -0.212 | 0.179 | 1.000 |
| VFLW vs HF | 0.074 | -0.097 | 0.244 | 0.845 |
| VFST vs HF | 0.173 | -0.022 | 0.369 | 0.115 |
| WF vs HF | 0.125 | -0.071 | 0.320 | 0.462 |
| RP vs OGF | -0.219 | -0.410 | -0.029 | **0.014** |
| VFLW vs OGF | -0.130 | -0.294 | 0.035 | 0.220 |
| VFST vs OGF | -0.030 | -0.220 | 0.160 | 0.999 |
| WF vs OGF | -0.078 | -0.269 | 0.112 | 0.872 |
| VFLW vs RP | 0.090 | -0.075 | 0.255 | 0.648 |
| VFST vs RP | 0.190 | -0.001 | 0.380 | 0.051 |
| WF vs RP | 0.141 | -0.049 | 0.331 | 0.283 |
| VFST vs VFLW | 0.100 | -0.065 | 0.265 | 0.529 |
| WF vs VFLW | 0.051 | -0.114 | 0.216 | 0.964 |
| WF vs VFST | -0.048 | -0.239 | 0.142 | 0.987 |

Signif. codes: 0 ‘***’ 0.001 ‘**’ 0.01 ‘*’ 0.05 ‘.’ 0.1 ‘ ’ 1

**Table SI 20:** Permutation test for homogeneity of multivariate dispersions for reptile species composition within seven land-use types.

|  | Df | Sum Sq | Mean Sq | F | N.Perm | Pr(>F) |  |
| --- | --- | --- | --- | --- | --- | --- | --- |
| Land-use management | 6 | 0.116 | 0.0194 | 1.870 | 999 | 0.088 | . |
| Residuals | 73 | 0.757 | 0.0104 |  |  |  |  |

Signif. codes: 0 ‘***’ 0.001 ‘**’ 0.01 ‘*’ 0.05 ‘.’ 0.1 ‘ ’ 1

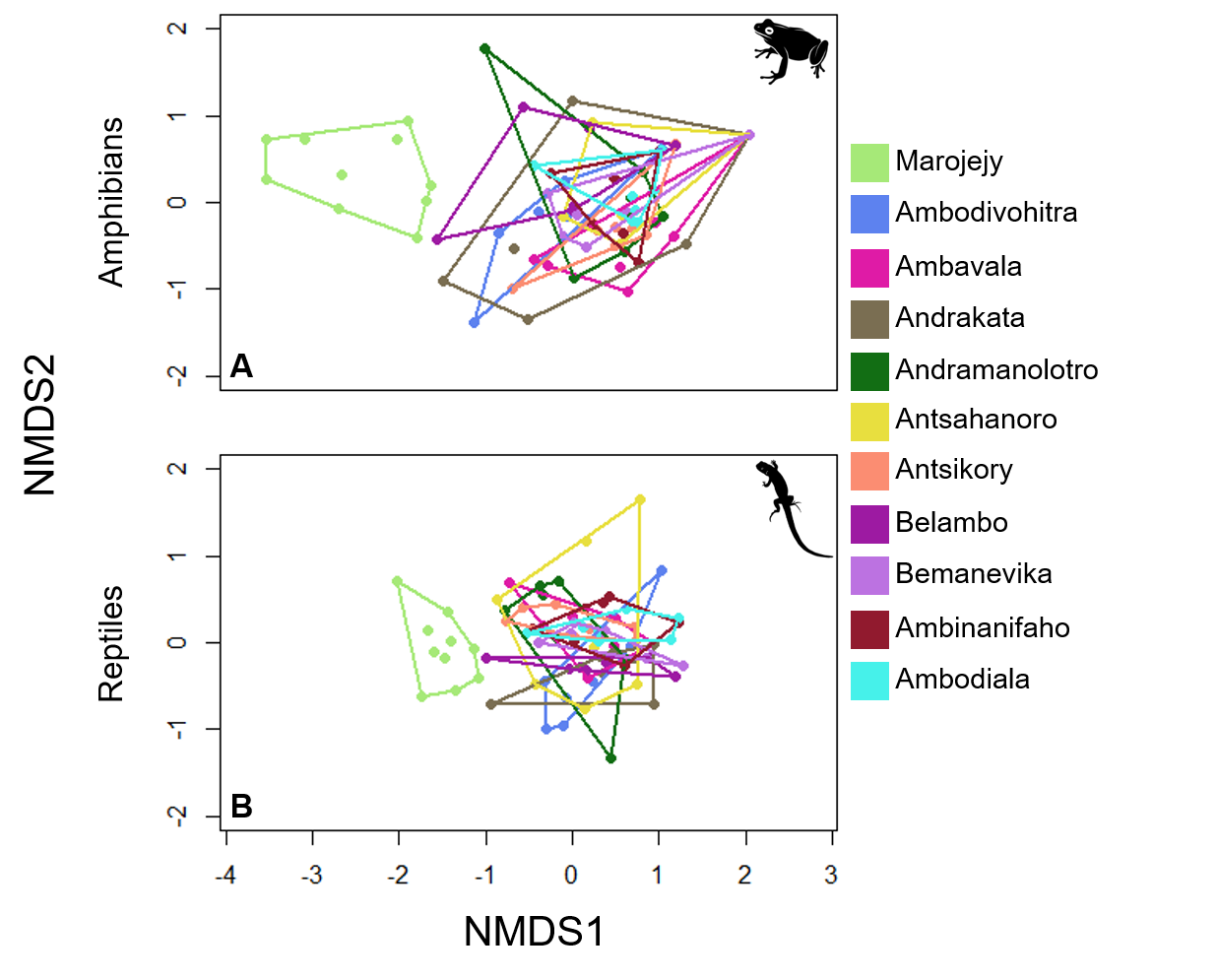
 **Figure SI 21:** Species composition of amphibian and reptile per village and in Marojejy National Park in north-eastern Madagascar. Non-metric dimensional scaling (NMDS) showing community dissimilarity of (A) amphibian and (B) reptile communities. Icon source: Free Icon Library (see reference below).

**Table SI 22**: Adonis test of community dissimilarity for amphibian species between villages and Marojejy National Park (**MJ**). Significant p-values (<0.05) are highlighted in bold.

|  | Df | SumsOfSqs | MeanSqs | F.Model | R2 | Pr(>F) | Sign. |
| --- | --- | --- | --- | --- | --- | --- | --- |
| Village and MJ | 10 | 11.928 | 1.193 | 4.849 | 0.416 | **<0.001** | *** |
| Residuals | 68 | 16.729 | 0.246 |  | 0.584 |  |  |
| Total | 78 | 28.657 |  |  | 1 |  |  |

Signif. codes: 0 ‘***’ 0.001 ‘**’ 0.01 ‘*’ 0.05 ‘.’ 0.1 ‘ ’ 1

**Table SI 23**: Adonis test of community dissimilarity for reptile species between villages and Marojejy National Park (**MJ**). Significant p-values (<0.05) are highlighted in bold.

|  | Df | SumsOfSqs | MeanSqs | F.Model | R2 | Pr(>F) | Sign. |
| --- | --- | --- | --- | --- | --- | --- | --- |
| Village and MJ | 10 | 11.004 | 1.100 | 4.398 | 0.389 | **<0.001** | *** |
| Residuals | 69 | 17.264 | 0.250 |  | 0.611 |  |  |
| Total | 79 | 28.268 |  |  | 1 |  |  |

Signif. codes: 0 ‘***’ 0.001 ‘**’ 0.01 ‘*’ 0.05 ‘.’ 0.1 ‘ ’ 1

**Table SI 24:** Pairwise-adonis for amphibian and reptile species composition between villages and Marojejy National Park. **MJ**: Marojejy National Park, **V2**: Ambavala, **V7**: Ambinanifaho, **V8**: Ambodiala, **V13**: Ambodivohitra, **V24**: Andrakata, **V25**: Andramanolotra, **V39**: Antsahanoro, **V40**: Antsikory, **V45**: Belambo, **V47**: Bemanevika

| number | pairs | Df | SumsOfSqs | F.Model | R2 | p.value | p.adjusted | sig |
| --- | --- | --- | --- | --- | --- | --- | --- | --- |
| **Amphibian** | | | | | | | | |
| 1 | MJ vs V13 | 1 | 3.420 | 12.916 | 0.463 | 0.001 | 0.055 |  |
| 2 | MJ vs V2 | 1 | 4.547 | 17.411 | 0.537 | 0.001 | 0.055 |  |
| 3 | MJ vs V24 | 1 | 3.599 | 11.402 | 0.432 | 0.001 | 0.055 |  |
| 4 | MJ vs V25 | 1 | 4.391 | 17.771 | 0.542 | 0.001 | 0.055 |  |
| 5 | MJ vs V39 | 1 | 4.733 | 19.481 | 0.565 | 0.001 | 0.055 |  |
| 6 | MJ vs V40 | 1 | 4.499 | 19.083 | 0.560 | 0.001 | 0.055 |  |
| 7 | MJ vs V45 | 1 | 2.865 | 10.158 | 0.420 | 0.001 | 0.055 |  |
| 8 | MJ vs V47 | 1 | 4.241 | 18.180 | 0.548 | 0.001 | 0.055 |  |
| 9 | MJ vs V7 | 1 | 4.692 | 20.618 | 0.579 | 0.001 | 0.055 |  |
| 10 | MJ vs V8 | 1 | 4.987 | 23.298 | 0.608 | 0.001 | 0.055 |  |
| 11 | V13 vs V2 | 1 | 0.683 | 2.517 | 0.173 | 0.037 | 1.000 |  |
| 12 | V13 vs V24 | 1 | 0.586 | 1.727 | 0.126 | 0.111 | 1.000 |  |
| 13 | V13 vs V25 | 1 | 0.725 | 2.856 | 0.192 | 0.012 | 0.660 |  |
| 14 | V13 vs V39 | 1 | 0.505 | 2.033 | 0.145 | 0.057 | 1.000 |  |
| 15 | V13 vs V40 | 1 | 0.572 | 2.388 | 0.166 | 0.028 | 1.000 |  |
| 16 | V13 vs V45 | 1 | 0.273 | 0.914 | 0.077 | 0.536 | 1.000 |  |
| 17 | V13 vs V47 | 1 | 0.207 | 0.876 | 0.068 | 0.504 | 1.000 |  |
| 18 | V13 vs V7 | 1 | 0.529 | 2.305 | 0.161 | 0.036 | 1.000 |  |
| 19 | V13 vs V8 | 1 | 0.721 | 3.393 | 0.220 | 0.019 | 1.000 |  |
| 20 | V2 vs V24 | 1 | 0.454 | 1.357 | 0.102 | 0.226 | 1.000 |  |
| 21 | V2 vs V25 | 1 | 0.354 | 1.422 | 0.106 | 0.175 | 1.000 |  |
| 22 | V2 vs V39 | 1 | 0.470 | 1.928 | 0.138 | 0.119 | 1.000 |  |
| 23 | V2 vs V40 | 1 | 0.236 | 1.003 | 0.077 | 0.437 | 1.000 |  |
| 24 | V2 vs V45 | 1 | 0.539 | 1.834 | 0.143 | 0.069 | 1.000 |  |
| 25 | V2 vs V47 | 1 | 0.345 | 1.489 | 0.110 | 0.237 | 1.000 |  |
| 26 | V2 vs V7 | 1 | 0.531 | 2.365 | 0.165 | 0.045 | 1.000 |  |
| 27 | V2 vs V8 | 1 | 0.456 | 2.195 | 0.155 | 0.075 | 1.000 |  |
| 28 | V24 vs V25 | 1 | 0.498 | 1.570 | 0.116 | 0.103 | 1.000 |  |
| 29 | V24 vs V39 | 1 | 0.469 | 1.504 | 0.111 | 0.191 | 1.000 |  |
| 30 | V24 vs V40 | 1 | 0.442 | 1.457 | 0.108 | 0.203 | 1.000 |  |
| 31 | V24 vs V45 | 1 | 0.276 | 0.749 | 0.064 | 0.686 | 1.000 |  |
| 32 | V24 vs V47 | 1 | 0.443 | 1.478 | 0.110 | 0.194 | 1.000 |  |
| 33 | V24 vs V7 | 1 | 0.553 | 1.887 | 0.136 | 0.067 | 1.000 |  |
| 34 | V24 vs V8 | 1 | 0.549 | 1.991 | 0.142 | 0.059 | 1.000 |  |
| 35 | V25 vs V39 | 1 | 0.159 | 0.702 | 0.055 | 0.720 | 1.000 |  |
| 36 | V25 vs V40 | 1 | 0.200 | 0.918 | 0.071 | 0.523 | 1.000 |  |
| 37 | V25 vs V45 | 1 | 0.504 | 1.836 | 0.143 | 0.043 | 1.000 |  |
| 38 | V25 vs V47 | 1 | 0.360 | 1.679 | 0.123 | 0.106 | 1.000 |  |
| 39 | V25 vs V7 | 1 | 0.165 | 0.796 | 0.062 | 0.611 | 1.000 |  |
| 40 | V25 vs V8 | 1 | 0.127 | 0.669 | 0.053 | 0.683 | 1.000 |  |
| 41 | V39 vs V40 | 1 | 0.168 | 0.793 | 0.062 | 0.564 | 1.000 |  |
| 42 | V39 vs V45 | 1 | 0.393 | 1.462 | 0.117 | 0.200 | 1.000 |  |
| 43 | V39 vs V47 | 1 | 0.200 | 0.954 | 0.074 | 0.464 | 1.000 |  |
| 44 | V39 vs V7 | 1 | 0.036 | 0.177 | 0.015 | 0.963 | 1.000 |  |
| 45 | V39 vs V8 | 1 | 0.135 | 0.730 | 0.057 | 0.617 | 1.000 |  |
| 46 | V40 vs V45 | 1 | 0.465 | 1.793 | 0.140 | 0.071 | 1.000 |  |
| 47 | V40 vs V47 | 1 | 0.250 | 1.251 | 0.094 | 0.355 | 1.000 |  |
| 48 | V40 vs V7 | 1 | 0.212 | 1.098 | 0.084 | 0.408 | 1.000 |  |
| 49 | V40 vs V8 | 1 | 0.206 | 1.169 | 0.089 | 0.316 | 1.000 |  |
| 50 | V45 vs V47 | 1 | 0.285 | 1.112 | 0.092 | 0.393 | 1.000 |  |
| 51 | V45 vs V7 | 1 | 0.365 | 1.471 | 0.118 | 0.187 | 1.000 |  |
| 52 | V45 vs V8 | 1 | 0.520 | 2.265 | 0.171 | 0.016 | 0.880 |  |
| 53 | V47 vs V7 | 1 | 0.230 | 1.212 | 0.092 | 0.337 | 1.000 |  |
| 54 | V47 vs V8 | 1 | 0.241 | 1.394 | 0.104 | 0.263 | 1.000 |  |
| 55 | V7 vs V8 | 1 | 0.165 | 0.996 | 0.077 | 0.448 | 1.000 |  |
| **Reptile** | | | | | | | | |
| 1 | MJ vs V13 | 1 | 2.576 | 9.040 | 0.376 | 0.001 | 0.055 |  |
| 2 | MJ vs V2 | 1 | 2.549 | 10.147 | 0.404 | 0.001 | 0.055 |  |
| 3 | MJ vs V24 | 1 | 3.643 | 15.321 | 0.505 | 0.001 | 0.055 |  |
| 4 | MJ vs V25 | 1 | 2.363 | 8.760 | 0.369 | 0.001 | 0.055 |  |
| 5 | MJ vs V39 | 1 | 3.459 | 12.837 | 0.461 | 0.001 | 0.055 |  |
| 6 | MJ vs V40 | 1 | 2.413 | 10.376 | 0.409 | 0.001 | 0.055 |  |
| 7 | MJ vs V45 | 1 | 3.003 | 11.949 | 0.443 | 0.001 | 0.055 |  |
| 8 | MJ vs V47 | 1 | 3.089 | 13.152 | 0.467 | 0.001 | 0.055 |  |
| 9 | MJ vs V7 | 1 | 3.490 | 14.044 | 0.484 | 0.001 | 0.055 |  |
| 10 | MJ vs V8 | 1 | 3.852 | 16.202 | 0.519 | 0.001 | 0.055 |  |
| 11 | V13 vs V2 | 1 | 0.418 | 1.440 | 0.107 | 0.189 | 1.000 |  |
| 12 | V13 vs V24 | 1 | 0.411 | 1.502 | 0.111 | 0.176 | 1.000 |  |
| 13 | V13 vs V25 | 1 | 0.432 | 1.378 | 0.103 | 0.215 | 1.000 |  |
| 14 | V13 vs V39 | 1 | 1.276 | 4.075 | 0.254 | 0.001 | 0.055 |  |
| 15 | V13 vs V40 | 1 | 0.697 | 2.608 | 0.179 | 0.010 | 0.550 |  |
| 16 | V13 vs V45 | 1 | 0.392 | 1.349 | 0.101 | 0.246 | 1.000 |  |
| 17 | V13 vs V47 | 1 | 0.464 | 1.719 | 0.125 | 0.113 | 1.000 |  |
| 18 | V13 vs V7 | 1 | 0.500 | 1.741 | 0.127 | 0.084 | 1.000 |  |
| 19 | V13 vs V8 | 1 | 0.666 | 2.436 | 0.169 | 0.023 | 1.000 |  |
| 20 | V2 vs V24 | 1 | 0.483 | 2.087 | 0.148 | 0.048 | 1.000 |  |
| 21 | V2 vs V25 | 1 | 0.313 | 1.154 | 0.088 | 0.276 | 1.000 |  |
| 22 | V2 vs V39 | 1 | 1.223 | 4.512 | 0.273 | 0.004 | 0.220 |  |
| 23 | V2 vs V40 | 1 | 0.096 | 0.427 | 0.034 | 0.903 | 1.000 |  |
| 24 | V2 vs V45 | 1 | 0.443 | 1.785 | 0.129 | 0.108 | 1.000 |  |
| 25 | V2 vs V47 | 1 | 0.122 | 0.536 | 0.043 | 0.851 | 1.000 |  |
| 26 | V2 vs V7 | 1 | 0.328 | 1.341 | 0.101 | 0.163 | 1.000 |  |
| 27 | V2 vs V8 | 1 | 0.286 | 1.234 | 0.093 | 0.243 | 1.000 |  |
| 28 | V24 vs V25 | 1 | 0.772 | 3.033 | 0.202 | 0.010 | 0.550 |  |
| 29 | V24 vs V39 | 1 | 1.282 | 5.043 | 0.296 | 0.002 | 0.110 |  |
| 30 | V24 vs V40 | 1 | 0.698 | 3.351 | 0.218 | 0.008 | 0.440 |  |
| 31 | V24 vs V45 | 1 | 0.328 | 1.415 | 0.105 | 0.195 | 1.000 |  |
| 32 | V24 vs V47 | 1 | 0.415 | 1.965 | 0.141 | 0.049 | 1.000 |  |
| 33 | V24 vs V7 | 1 | 0.503 | 2.206 | 0.155 | 0.025 | 1.000 |  |
| 34 | V24 vs V8 | 1 | 0.520 | 2.421 | 0.168 | 0.014 | 0.770 |  |
| 35 | V25 vs V39 | 1 | 1.232 | 4.189 | 0.259 | 0.001 | 0.055 |  |
| 36 | V25 vs V40 | 1 | 0.365 | 1.471 | 0.109 | 0.133 | 1.000 |  |
| 37 | V25 vs V45 | 1 | 0.729 | 2.683 | 0.183 | 0.006 | 0.330 |  |
| 38 | V25 vs V47 | 1 | 0.503 | 2.003 | 0.143 | 0.032 | 1.000 |  |
| 39 | V25 vs V7 | 1 | 0.556 | 2.075 | 0.147 | 0.019 | 1.000 |  |
| 40 | V25 vs V8 | 1 | 0.643 | 2.526 | 0.174 | 0.016 | 0.880 |  |
| 41 | V39 vs V40 | 1 | 1.528 | 6.167 | 0.339 | 0.001 | 0.055 |  |
| 42 | V39 vs V45 | 1 | 0.990 | 3.652 | 0.233 | 0.010 | 0.550 |  |
| 43 | V39 vs V47 | 1 | 1.033 | 4.122 | 0.256 | 0.001 | 0.055 |  |
| 44 | V39 vs V7 | 1 | 0.721 | 2.693 | 0.183 | 0.009 | 0.495 |  |
| 45 | V39 vs V8 | 1 | 1.017 | 3.999 | 0.250 | 0.003 | 0.165 |  |
| 46 | V40 vs V45 | 1 | 0.581 | 2.582 | 0.177 | 0.020 | 1.000 |  |
| 47 | V40 vs V47 | 1 | 0.233 | 1.139 | 0.087 | 0.326 | 1.000 |  |
| 48 | V40 vs V7 | 1 | 0.411 | 1.856 | 0.134 | 0.058 | 1.000 |  |
| 49 | V40 vs V8 | 1 | 0.337 | 1.618 | 0.119 | 0.124 | 1.000 |  |
| 50 | V45 vs V47 | 1 | 0.295 | 1.296 | 0.097 | 0.257 | 1.000 |  |
| 51 | V45 vs V7 | 1 | 0.380 | 1.553 | 0.115 | 0.106 | 1.000 |  |
| 52 | V45 vs V8 | 1 | 0.564 | 2.437 | 0.169 | 0.015 | 0.825 |  |
| 53 | V47 vs V7 | 1 | 0.229 | 1.019 | 0.078 | 0.404 | 1.000 |  |
| 54 | V47 vs V8 | 1 | 0.260 | 1.234 | 0.093 | 0.245 | 1.000 |  |
| 55 | V7 vs V8 | 1 | 0.245 | 1.073 | 0.082 | 0.359 | 1.000 |  |

Signif. codes: 0 ‘***’ 0.001 ‘**’ 0.01 ‘*’ 0.05 ‘.’ 0.1 ‘ ’ 1

**Table SI 25:** Permutation test for homogeneity of multivariate dispersions for amphibian species composition within villages and Marojejy National Park **(MJ)**.

|  | Df | Sum Sq | Mean Sq | F | N.Perm | Pr(>F) |
| --- | --- | --- | --- | --- | --- | --- |
| Village and MJ | 11 | 0.319 | 0.029 | 1.254 | 999 | 0.283 |
| Residuals | 67 | 1.547 | 0.023 |  |  |  |

Signif. codes: 0 ‘***’ 0.001 ‘**’ 0.01 ‘*’ 0.05 ‘.’ 0.1 ‘ ’ 1

**Table SI 26:** Permutation test for homogeneity of multivariate dispersions for reptile species composition within villages and Marojejy National Park **(MJ)**.

|  | Df | Sum Sq | Mean Sq | F | N.Perm | Pr(>F) |
| --- | --- | --- | --- | --- | --- | --- |
| Village and MJ | 11 | 0.090 | 0.008 | 0.835 | 999 | 0.626 |
| Residuals | 68 | 0.669 | 0.010 |  |  |  |

Signif. codes: 0 ‘***’ 0.001 ‘**’ 0.01 ‘*’ 0.05 ‘.’ 0.1 ‘ ’ 1

Reference

https://icon-library.com/115500-200.html>Amphibian,Frog,Vertebrate,Hyla,Tree frog,Toad,Poison dart frog,Tree frog,True frog,Clip art,Wood Frog,Shrub frog,Eleutherodactylus,Bufo,Phyllobates,Illustration # 185503

https://icon-library.com/icon/reptile-icon-20.html.html>Reptile Icon # 8802
